# Rapid Prototyping of Wireframe Scaffolded DNA Origami using ATHENA

**DOI:** 10.1101/2020.02.09.940320

**Authors:** Hyungmin Jun, Xiao Wang, William P. Bricker, Steve Jackson, Mark Bathe

**Affiliations:** Department of Biological Engineering, Massachusetts Institute of Technology, Cambridge, MA 02139, USA

**Author notes:** William P. Bricker, Department of Chemical and Biological Engineering, University of New Mexico, Albuquerque, NM 87131, USA.

## Abstract

Wireframe DNA origami assemblies can now be programmed automatically from the “top-down” using simple wireframe target geometries, or meshes, in 2D and 3D geometries using either rigid, six-helix bundle (6HB) or more compliant, two-helix bundle (2HB or DX) edges. While these assemblies have numerous applications in nanoscale materials fabrication due to their nanoscale spatial addressability and high degree of customization, no easy-to-use graphical user interface software yet exists to deploy these algorithmic approaches within a single, stand-alone interface. Here, we present ATHENA, an open-source software package with a graphical user interface that automatically renders single-stranded DNA scaffold routing and staple strand sequences for any target wireframe DNA origami in 2D or 3D using 2HB or 6HB edges. ATHENA enables external editing of sequences using the popular tool caDNAno, demonstrated here using asymmetric nanoscale positioning of gold nanoparticles, as well as atomic-level models for molecular dynamics, coarse-grained dynamics, or other computational chemistry simulation approaches. We anticipate ATHENA will significantly reduce the barrier for non-specialists to perform wireframe DNA origami sequence design and fabrication for custom applications in materials science, nanotechnology, therapeutics, and other areas.

## INTRODUCTION

Structural DNA nanotechnology was conceived in Ned Seeman’s pioneering work (1) in which he postulated that synthetic DNA could be used to program synthetic materials with prescribed nanometer-scale structural features. The use of synthetic oligonucleotides by Seeman resulted in extended, crystalline-like self-assembled DNA-based materials from objects of finite extent. Over two decades later, Paul Rothemund introduced the concept of scaffolded DNA origami (2) based on Seeman’s original design rules but now applied to the long, single-strand DNA genome of M13mp18 that he used to template dozens to hundreds of shorter, complementary synthetic DNA strands that self-assemble (or “fold”, hence “origami”) to form a single, discrete product with high yield. While the M13mp18 phage genome is still the most common scaffold used for this purpose, Rothemund’s approach applies generally to any scaffold length and sequence, which may be produced enzymatically (3, 4) or bacterially (5, 6). Soon after, Douglas et al., applied Rothemund’s approach to self-assemble 3D assemblies (7) based on similar design rules, and also released the widely used software tool caDNAno (8) to assist in the manual design of this class of scaffolded DNA origami in which DNA duplexes are arranged on parallel honeycomb or square lattices, also termed “bricklike” origami. While caDNAno has proven extremely useful for the manual scaffold routing and semi-automated sequence design of complementary staples used to self-assemble or fold target shapes, it has limited utility for curved and bent assemblies that require manual insertions and/or deletions in order to induce bend or twist (9, 10), or a new class of wireframe scaffolded DNA origami assemblies that render complex 2D and 3D target geometries based on simple wireframe “meshes” (3, 11–18).

Wireframe DNA origami design using double crossover (DX) or two-helix bundle (2HB) edges was first realized by Yan et al., (11) with the self-assembly of tiles, which was later generalized to 2D and 3D scaffolded DNA origami by Zhang et al., (13), and to incorporate single duplex edges by Benson et al., (12). In 2016, Veneziano et al., (3) demonstrated that arbitrary 3D wireframe geometries based on DX edges alone could be designed fully automatically based on target geometry, using DAEDALUS. In 2019, Jun et al. demonstrated an automatic design procedure for complex 2D wireframe DNA origami without any restrictions on edge length or geometric symmetry based on DX edges, called PERDIX (15). Soon thereafter, a similar design principle was applied for generating six-helix bundle (6HB) edge 3D assemblies (TALOS)(16) and 2D assemblies (METIS)(17) by Jun et al. The 6HB edge-based 2D and 3D assemblies showed significantly enhanced mechanical stiffness with respect to DX-edges, highlighting the potential of using wireframe DNA origami for constructing complex nanoscale materials facilitated by automatic design procedures (17).

While PERDIX (15), METIS (17), DAEDALUS (3), and TALOS (16) are versatile for the rendering of arbitrary 2D and 3D wireframe objects using scaffolded DNA origami, and are offered both online as free tools and downloadable open source software packages, they are still limited by their lack of a Graphical User Interface (GUI) to easily apply them to wireframe target geometries. And while caDNAno in principle offers the ability to perform wireframe scaffolded DNA origami design, in practice it is limited to experts with advanced knowledge of scaffold routing and staple sequence design rules and may require many hours for the sequence design of each target wireframe origami.

To enable practical and widely accessible fully automated sequence design of wireframe DNA scaffolded origami assemblies, here we introduce ATHENA, a GUI that integrates 2D and 3D target wireframe geometry file input together with application of fully automated sequence design and visualization. In addition to sequence design, ATHENA produces output files including all-atom structures in Protein Data Bank (PDB) (19) format for molecular visualization using tools such as Visual Molecular Dynamics (VMD) (20) or UCSF Chimera (21), all-atom molecular dynamics simulation, or coarse-grained simulation using tools such as oxDNA (22, 23), as well as caDNAno files for editing or modifying sequence designs for DNA origami functionalization or other purposes, and complete sequence files for ordering staple oligonucleotide strands required for fabrication via one-pot self-assembly.

## MATERIAL AND METHODS

### GUI implementation

ATHENA is an open source GUI software application (https://github.com/lcbb/athena) that performs fully automated sequence design of 2D or 3D wireframe scaffold DNA origami objects based on DX- or 6HB-based edges. ATHENA was implemented in Python using the Qt5 libraries, providing native support for both Windows and Mac operating systems. The back-end software packages such as PERDIX, DAEDALUS, METIS, and TALOS are embedded as binaries for executing jobs.

### PDB generation

The PDB generation software in ATHENA utilizes the nucleic-acid base-level nodes that are output from the routing procedure, and these nodes include information on the sequence, routing, and position of each nucleic acid base. The first step in the PDB generation is to route the base-level node information into sequential nucleic acid strands appropriate for an all-atom model, which is accomplished by a searching algorithm since each base is mapped to the upstream, downstream, and paired bases in the model. Next, the all-atom model is built base-by-base and strand-by-strand by transforming the coordinates of a reference average B-form nucleic acid base structure onto the node-level positions. The all-atom nucleic acid structures used are from the 3DNA parameter set (24), where the coordinates are based on average B-form DNA structures from Olson, et al. (25). Several ProDy coordinate transformation functions are utilized during PDB generation (26). Single-stranded nucleic acid regions are not included in the node-level routing, so the unpaired coordinates are interpolated from the nearest upstream and downstream base-pairs using a cubic Bézier function, providing a smooth path from arbitrary base-pair coordinates.

The standard PDB file format (19) has several longstanding limitations for large atomic structures, including limitations on the number of separate chains or nucleic acid strands (62, case-sensitive alphanumeric), the number of total atoms (99,999), the number of residues or nucleic acid bases (9,999), and the spatial dimensions {-999.999, 9999.999} in Ångstroms. This PDB generation software utilizes workarounds for some of these limitations. The atom numbering scheme above index 99,999 utilizes a hybrid base-36 encoding scheme where the first character is case-sensitive alphabetical and the following four characters are base-36 alphanumeric, in theory allowing for >87 million total atoms. The alphabetical first character allows any parser to recognize the switch from base-10 to hybrid base-36 encoding. The residue numbering scheme above index 9,999 similarly allows for >2.4 million total residues using the same hybrid base-36 encoding. For larger atomic structures, in particular with spatial dimensions exceeding the standard PDB limitations, the PDBx/mmCIF file format (27) could be utilized, but this is left for future work.

### Materials

DNA origami staple strands were purchased in 96-well plate format from Integrated DNA Technologies, Inc. at 25-nmole synthesis scale. The staple strands were purified by standard desalting and calibrated to 200 μM based on full yield. Staple strands were mixed in equal volume from the corresponding wells and used directly for DNA origami folding without further purification. 5’ Thiol Modifier (C6 S-S) modified DNA strand was purchased from Integrated DNA Technologies, Inc. at 100-nmole synthesis scale with standard desalting. Nuclease Free Water was purchased from Integrated DNA Technologies, Inc. The 7,249-nt DNA scaffold (M13mp18) was purchased from Guild BioSciences at a concentration of 100 nM. 10x TAE buffer was purchased from Alfa Aesar. Magnesium acetate tetrahydrate (molecular biology grade) was purchased from MilliporeSigma. 1x TAE buffer with 12.5 mM Mg(OAc)_2_ was prepared with 10x TAE buffer and Magnesium acetate tetrahydrate. 5nm OligoREADY Gold Nanoparticle Conjugation Kit was purchased from Cytodiagnostics Inc. Pierce DTT (Dithiothreitol) was purchased from Thermo Scientific, and illustra NAP-5 columns were purchased from GE Healthcare Life Sciences.

### Origami Self-assembly

All pentagonal DNA origami objects were folded with the same protocol. 5 nM of DNA scaffold was mixed with 20 equiv corresponding staples strands in 1x TAE buffer with 12.5 mM Mg(OAc)_2_, the final volume of the self-assembly solution was 100 μl. The mixture buffer solution was annealed in a PCR thermocycler: 95 °C for 2 min, 70 °C to 45 °C at a rate of 0.5 °C per 20 min, and 45 °C to 20°C at a rate of 0.5 °C per 10 min. The annealed solution was validated by 1.5% Agarose gel in 1x TAE buffer with 12.5 mM Mg(OAc)2 and and 1x SybrSafe. Gels were run at 60 V and subsequently imaged under blue light. The annealed solution was diluted into 500 μl with 1x TAE buffer with 12.5 mM Mg(OAc)_2_, and the extra staple strands were removed with MWCO = 100 kDa spin filter concentration columns. The purified DNA origami solution was adjusted to desired concentrations (5 nM) for AFM and TEM imaging.

### Preparation of DNA-gold NP Conjugate Modified DNA Origami

The 5’ Thiol Modifier modified DNA strand (50 μM) was reduced by DTT (0.1 M) in 0.15 M sodium phosphate buffer (pH 8.5) for 2 hours at room temperature. The reaction solution was then purified with Nap-5 column to remove small molecules from 5’ thiol-DNA strand. The purified 5’ thiol-DNA strand was adjusted to 25 μM in nuclease free water based on the OD_260nm_. One vial of lyophilized OligoREADY™ 5nm gold nanoparticle was resuspended in 740 µl of nuclease free H_2_O. Add 160 µl of purified 5’ thiol-DNA strand (25 μM) and 100 µl of 1M NaCl to the gold NP suspension. The mixture was incubated at room temperature for 2 hours. The excess DNA strand was subsequently removed from MWCO = 100 kDa spin filter concentration columns, and the DNA-gold NP conjugate was concentrated in the meantime. The concentration of DNA-gold NP conjugate was determined by OD_520nm_.

The DNA-gold NP conjugate was added to purified DNA origami solution (20 nM) in a ratio of 5: 1 (gold NP: sites of modification on origami), and the mixtures were incubated in 1x TAE buffer with 12.5 mM Mg(OAc)_2_ at room temperature overnight.

### AFM and TEM Imaging

AFM imaging was performed in “ScanAsyst mode in fluid” (Dimension FastScan, Bruker Corporation) with ScanAsyst-Fluid+ or SNL-10 tips (Bruker Inc.). Two microliters of sample (5 nM) were deposited onto freshly cleaved mica (Ted Pella Inc.), and 0.5 to 1.0 μl of NiCl2 at a concentration of 100 mM were added to the samples to fix the origami nanostructures on the mica surface. After waiting for approximately 30 s for sample adsorption to mica, 80 μl of 1x TAE/Mg2+ buffer was added to the samples, and an extra 40 μl of the same buffer was deposited onto the AFM tip. For TEM imaging, 5 uL of DNA origami solution (5 nM) was deposited onto fresh glow discharged carbon film with copper grids (CF200H-CU; Electron Microscopy Sciences Inc., Hatfeld, PA), and the sample was then allowed to absorb onto the surface for 30s. After the sample solution was blotted from the grid using Whatman 42 filter paper, the grid was placed on 5 uL of freshly prepared 2% uranyl-formate with 25 mM NaOH for 10s. The remaining stain solution on the grid was blotted away using Whatman 42 filter paper and dried under house vacuum prior to imaging. The sample was imaged on a Technai FEI with a Gatan camera.

## RESULTS AND DISCUSSION

ATHENA provides fully automated sequence design of 2D or 3D wireframe scaffold DNA origami objects based uniformly either on rigid 6HB or more compliant 2HB edges (Figure 1 and Supplementary Note 1). 2D and 3D target geometries must be specified using a polygonal surface mesh and, in 3D, each edge of every polygonal surface should be one of the edges of a neighboring surface. These are provided manually or through an ASCII file format that define the polygonal mesh, such as the Polygon File Format (PLY), STereoLithography (STL), or Virtual Reality Modeling Language (WRL) using any number of standard CAD programs. A PLY file is used as input to Athena because of its simplicity and broad use within CAD-based design. Conversion from STL to PLY filetypes may be performed using open source tools and online convertors. Because scaffold routing and staple design are based on PLY files, it is essential that every vertex listed in the file pertain to at least one face, since otherwise there is no way of routing the ssDNA scaffold through the entire target object (16). Once input, ATHENA offers the ability to visualize the target 2D or 3D wireframe object using surface shading and/or wireframe edges, in default colors that may be altered using custom options (Figure 1(a); i). Zooming, rotation, and translation may each be selected as standard mouse options, as well as Perspective versus Orthographic views (Figure 1(a); i). ATHENA also provides pre-defined target geometries of 37 for 2D and 55 for 3D (Figure 1(a); ii and Supplementary Notes 1 and 2).

**Figure 1.**
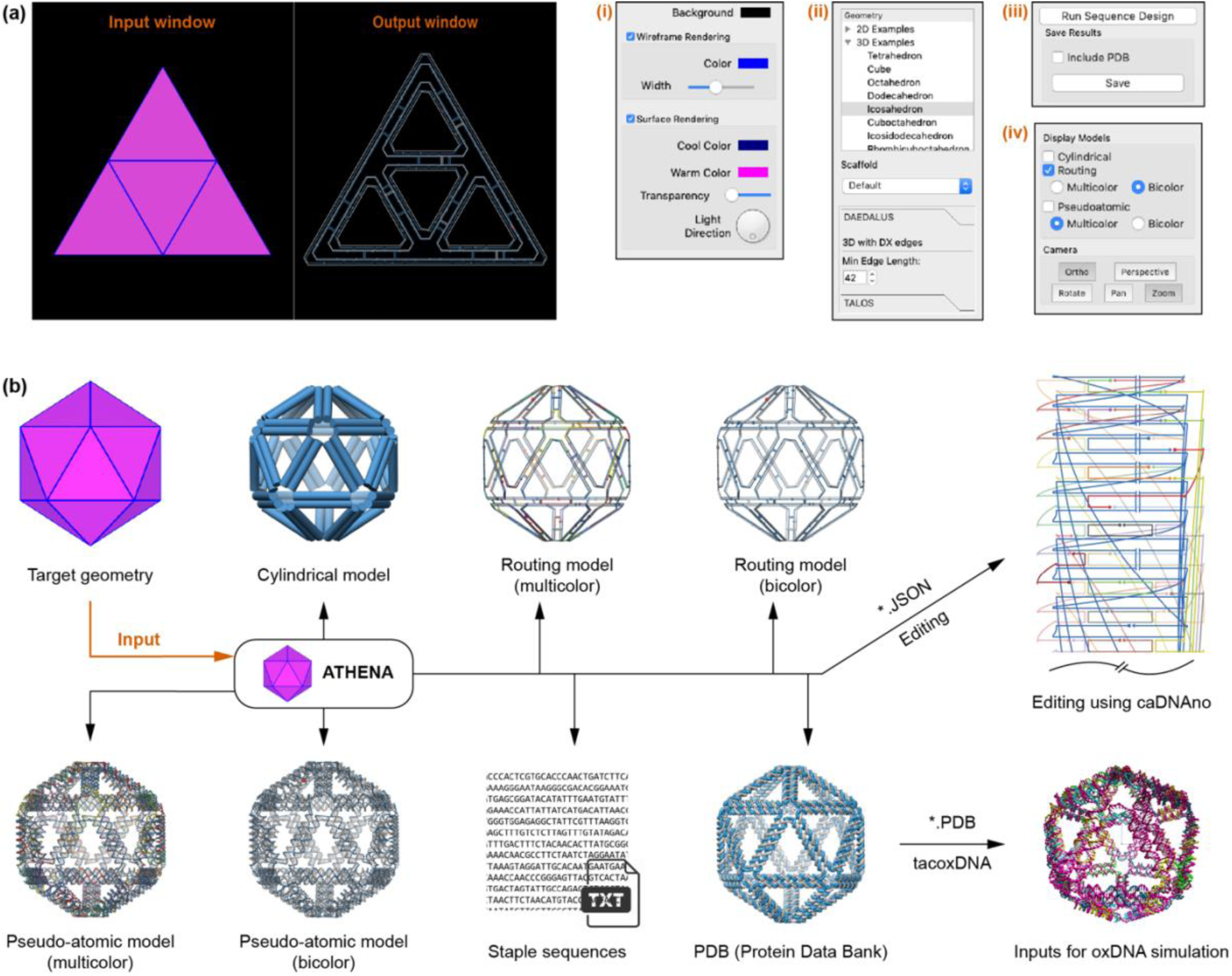
Interface and design outputs of ATHENA. (**a**) Screenshot of graphical user interface that has two windows for rendering the target geometry (input window) and outputs (output window) such as cylindrical, routing, and pseudo-atomic model. Additional four panels are to control options; rendering colour scheme, target geometry, scaffold sequence, edge length, edge type, camera control, and outputs. (**b**) Based on the target geometry, ATHENA routes a single-stranded scaffold throughout the entire geometry and generates several outputs; cylindrical model, routing model, pseudo-atomic model, text file for staple sequences, JSON for caDNAno, and PDB for molecular dynamic simulations.

ATHENA uses M13mp18 as a default scaffold sequence for required lengths less than or equal to 7,249-nt, a Lambda phage sequence if greater than 7,250-nt and less than or equal to 48,502-nt, and a random sequence if greater than 48,503-nt. User-defined scaffold sequences can also be imported using a text file (Figure 1(a); ii). ATHENA has the option to choose the edge type; DNA double-crossover (DX or 2HB) or six-helix bundle (6HB) that consists of every edge of the 2D or 3D wireframe objects (Figure 1(a); ii). Then, fully automated scaffold and staple sequence design can be performed using either DX- (3, 15) or 6HB-edge (16, 17) motifs with either the default, M13 ssDNA scaffold, or a custom scaffold of length and sequence defined by the user (Figure 1(a); iii). In addition, the minimum edge length is assigned to the shortest edge, which is then used to scale all other edges, specifying from 42-bp (13.9 nm) to 210-bp (71.1 nm) edge-lengths in the design, which may range from 20 nm to 200 nm for 2D and from 20 nm to 100 nm for 3D when using the M13mp18 ssDNA scaffold (7,249-nt).

Once the sequence design procedure in ATHENA is completed, the cylindrical representation is displayed overlapping with the target geometry (Figure 1(b)). In the cylindrical model, each edge of the wireframe structure is rendered using a cylinder (2 nm diameter) that represents a DNA double helix. Strand routing and the helicity of DNA can be displayed using the routing and pseudo-atomic model options (Figure 1(a); iv). For the routing model, each strand, including the scaffold and staples, is approximated by a vector representing the direction of the DNA strand (Figure 1(b)). More detailed output with the double-helical DNA can be displayed in the pseudo-atomic model constructed by spheres and lines representing nucleotides and the backbone of DNA, respectively. For easier identification of the scaffold and individual staples, two colour schemes with multiple colours are built for the routing and pseudo-atomic models. The resulting sequence outputs can also be exported (Figure 1(a); iii) with several files; a Comma Separated Values (CSV) spreadsheet containing staple sequences, a PDB all-atom model, and JavaScript Object Notation (JSON) for caDNAno (Figure 1(b)). The tacoxDNA (28) webserver can be used to convert the PDB file to the appropriate inputs for performing coarse-grained simulations with oxDNA (22, 23). The JSON file can be imported into caDNAno (8) for manual base and oligo editing for functionalization, for example, editing sequences, extending strands, deleting and adding nucleotides, and changing the position for crossovers and nicks (Figure S1).

Based only on a target geometry, scaffold sequence, and edge type (DX or 6HB), ATHENA performs automated scaffold routing and staple sequence design, and generates the required staple strands needed to experimentally fold the structure. PERDIX performs fully automated scaffold routing and staple sequence design for any free-form 2D geometry using exclusively DX-based edges, whereas METIS designs any 2D geometry using mechanically stiffer honeycomb or 6HB edges. DAEDALUS solves the scaffold routing and staple design problem fully automatically for any 3D polyhedral surface using solely DX-based edges, whereas TALOS renders any 3D polyhedral surface using mechanically stiffer honeycomb edges, thereby also requiring greater scaffold length for the same particle geometry and size. TALOS additionally offers the ability to utilize every crossover possible between neighboring 6HB edge duplexes (16), which should offer enhanced mechanical and enzymatic integrity compared with the minimal number of single crossovers utilized between any two edges in the original sequence designs of Tian et al. (29).

We tested the ability of ATHENA to generate high-quality wireframe DNA origami structures, which also allows users to further functionalize such structures with other materials conveniently. Here, we generated the staple strands sequences of five 6HB-based pentagonal objects (Figure 2 and Tables S1-S7) with different edge-lengths from 42-bp (13.94 nm) to 210-bp (71.06 nm) with ATHENA. Transmission electron microscopy (TEM) and atomic force microscopy (AFM) confirmed the successful assembly of target structures as indicated by the accurate vertex angles and the high yield of proper formation of these structures (Figures S2-S11). Users can modify these structures based on the routing and pseudo-atomic model generated by ATHENA, which enables the user to identify the position of a particular modification (nick or overhang position). Each staple strand was labelled with the same color in both the pseudo-atomic model and caDNAno file, for convenience in identifying the corresponding staple strands in the caDNAno file for modifications. To demonstrate the addressability of this well-controlled scaffolding material and editing approach, we modified one of the pentagonal origami structures (210-bp edge length) for gold nanoparticle attachment (Figure 3). Following the procedure described in Figure S12, we modified staple strands around the vertex of this pentagonal structure for positioning gold nanoparticles. The handles for DNA-gold NP conjugates were placed at either three or all five vertices of the pentagonal structure, and the handles from the adjacent edges were designed to fix one gold nanoparticle in the vertex. (Figure 3(c) and Figures S13-S15). TEM images showed that the gold nanoparticles were successfully placed at the prescribed positions in the origami structure, which alternatively could be used to program any number of inorganic or organic molecules, in both 2D and 3D, as commonly performed in the DNA origami field (29).

**Figure 2.**
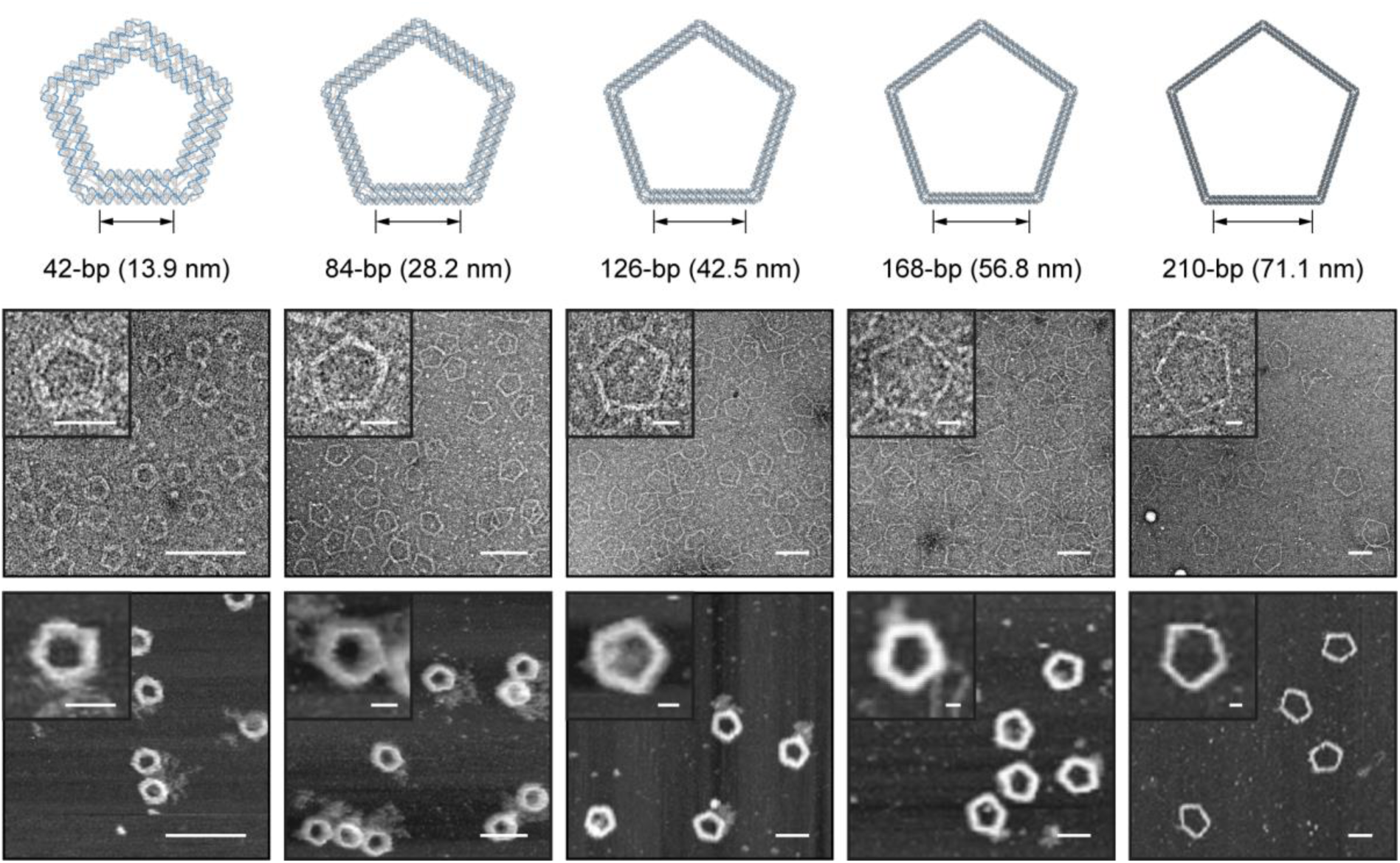
Designing 6HB pentagonal DNA origami objects with variable edge lengths. TEM and AFM images for variable edge lengths of 42-, 84-, 126-, 168-, and 210-bp DNA pentagonal objects. Scale bars, 25 nm and 100 nm (zoom-in and zoom-out TEM and AFM images, respectively).

**Figure 3.**
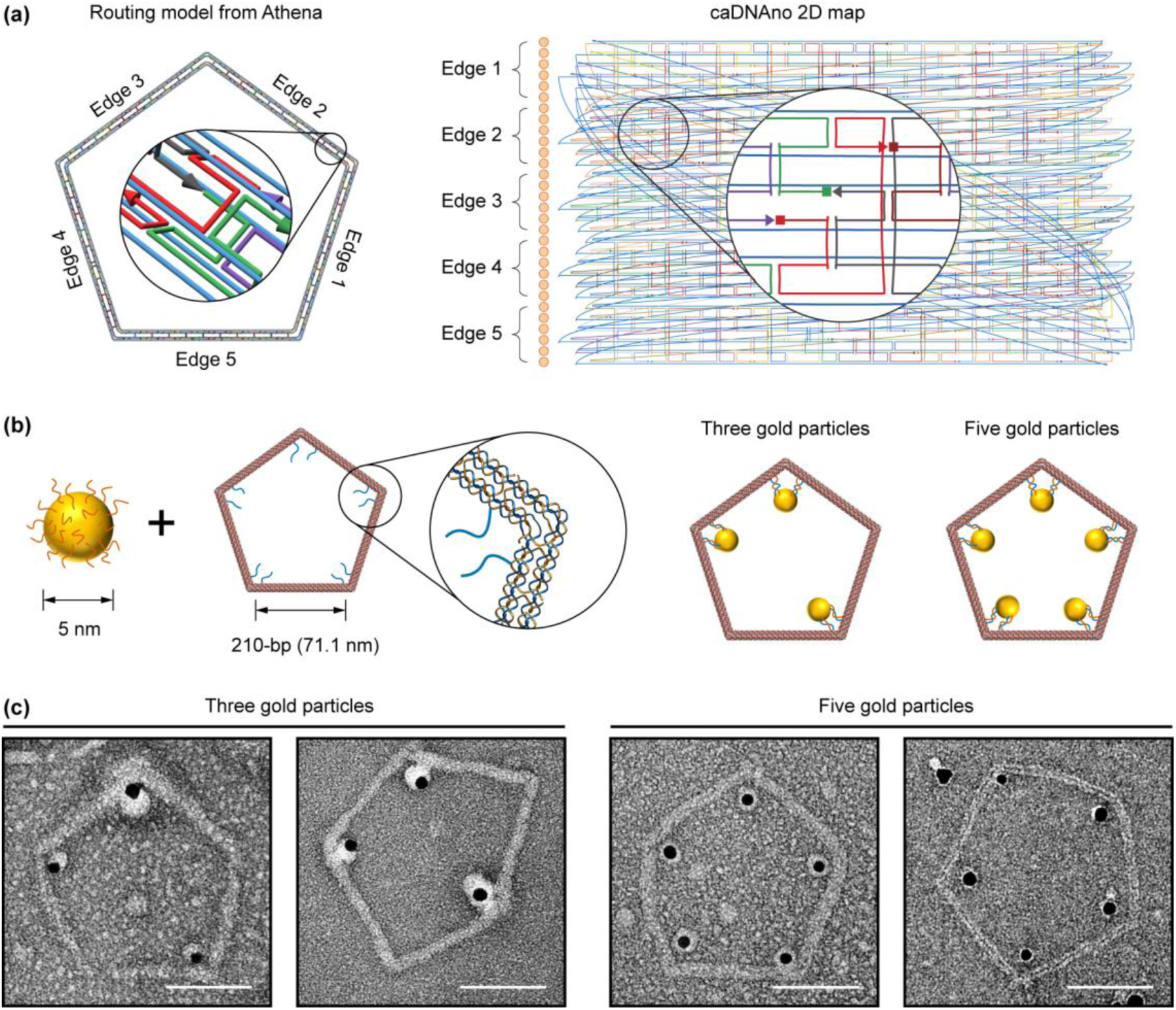
Organizing gold nanoparticles on the pentagonal DNA origami. (**a**) Routing model and caDNAno representation for 210-bp pentagonal DNA origami design from Athena. (**b**) Diagrams showing the attachment of nanoparticles at every three or five corners. (**c**) Transmission electron micrographs of 2D pentagonal DNA origami of organized gold nanoparticles. Scale bars, 50 nm.

## SUPPLEMENTARY DATA

Supplementary Data are available online.

