## Supplementary Information for "Rapid Prototyping of Wireframe Scaffolded DNA Origami using ATHENA"

**Rapid Prototyping of Structured DNA Assemblies using ATHENA**

**Supplementary Note 1.** ATHENA options reference.

**Main menu - File**

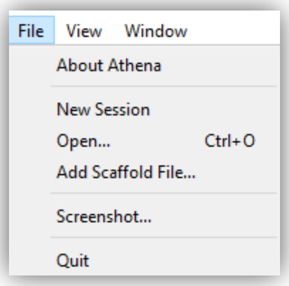


| Name | Function |
| --- | --- |
| About Athena | Software information |
| New Session | Make new session |
| Open | Open custom PLY file |
| Add Scaffold File | Open the text file for custom scaffold sequences |
| Screenshot | Captures the working window and export it to the image file |
| Quit | Quit ATHENA |

**Main menu - View**

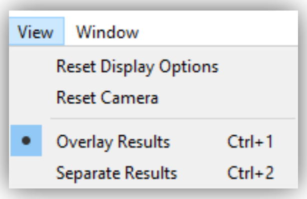


| Name | Function |
| --- | --- |
| Reset Display | Reset display setting as default |
| Reset Camera | Reset camera setting as default |
| Overlay Results | One window; output rendering is overlapped on the target geometry |

| | |
|---|---|
| Separate Results | Two windows; one for the target geometry and another for output rendering |

**Main menu - Window**

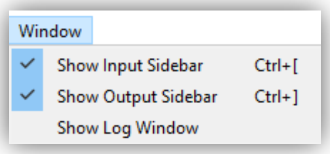


| | |
|---|---|
| Show Input Sidebar | Show / hide input sidebar |
| Show Output Sidebar | Show / hide output sidebar |
| Show Log Window | Log window displays design parameters generated by Athena when it runs |

**Main menu - Window**

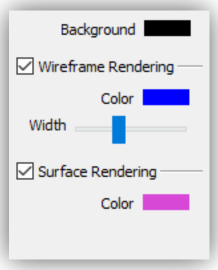


| | |
|---|---|
| Background | Change the background (black as default) |
| Wireframe Rendering | Change the color or width of the edge of the target geometry (blue as default) |
| Surface Rendering | Change the color for the surface of the target geometry (magenta as default) |

**Display Models**

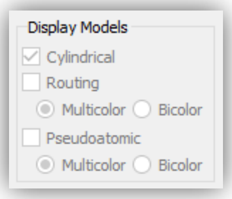


| | |
|---|---|
| Cylindrical | Cylindrical model representation<br>Each cylinder represents the DNA duplex |
| Routing | |
| Pseudoatomic | |

**Camera**

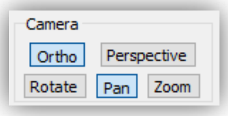


| | |
|---|---|
| Ortho | Orthographic projection |
| Perspective | perspective projection |
| Rotate | Rotate the object in the view window |
| Pan | Translate the object in the view window |
| Zoom | Zoom in/out the object in the view window |

**Geometry**

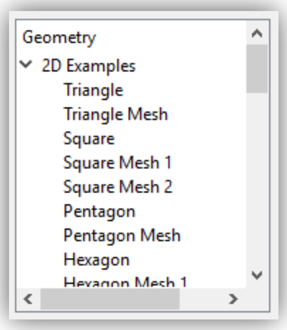


| | |
|---|---|
| 2D Examples | 37 pre-defined 2D target geometries |
| 3D Examples | 55 pre-defined 3D target geometries |
| Loaded Files | Display loaded target geometry by user |

##### Scaffold

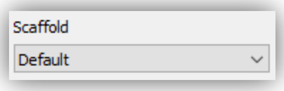


| | |
|---|---|
| Default | M13mp18 as a default scaffold sequence for required length less than or equal to 7,249-nt, a Lambda phage sequence if greater than 7,250-nt and less or equal to 48,502-nt, and a random sequence if greater than 48,503-nt |

##### PERDIX - DX-based 2D wireframe origami

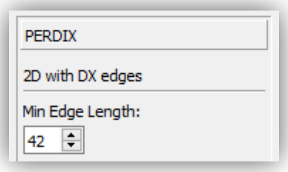


| | |
|---|---|
| Min Edge Length | The minimum edge length (from 42-bp to 126-bp) is assigned to the shortest edge and the other edges are scaled and rounded appropriately |

##### METIS - 6HB-based 2D wireframe origami

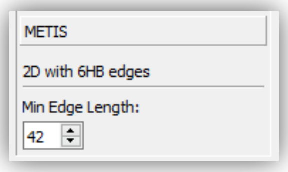


| | |
|---|---|
| Min Edge Length | The minimum edge length (from 42-bp to 126-bp) is assigned to the shortest edge and the other edges are scaled and rounded appropriately |

##### DAEDALUS - DX-based 3D wireframe origami

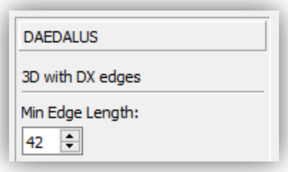


| | |
|---|---|
| Min Edge Length | The minimum edge length (from 42-bp to 126-bp) is assigned to the shortest edge and the other edges are scaled and rounded appropriately |

##### TALOS - 6HB-based 3D wireframe origami

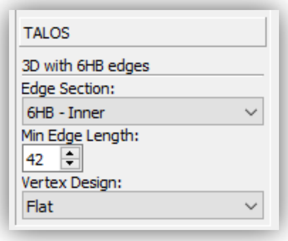


| | |
|---|---|
| Edge Section | Inner / Middle connections |
| Min Edge Length | The minimum edge length (from 42-bp to 126-bp) is assigned to the shortest edge and the other edges are scaled and rounded appropriately |
| Vertex Design | Flat vs mitered vertex design |

##### Run

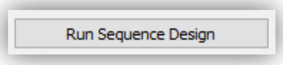


| | |
|---|---|
| Sequence Design | Run to sequence design |

##### Save Results

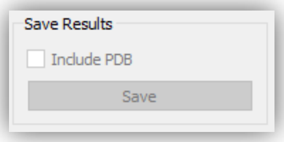


| | |
|---|---|
| Include PDB | Save outputs with PDB file |
| Save | Save outputs as files |

**Supplementary Note 2.** 37 pre-defined 2D target geometries.

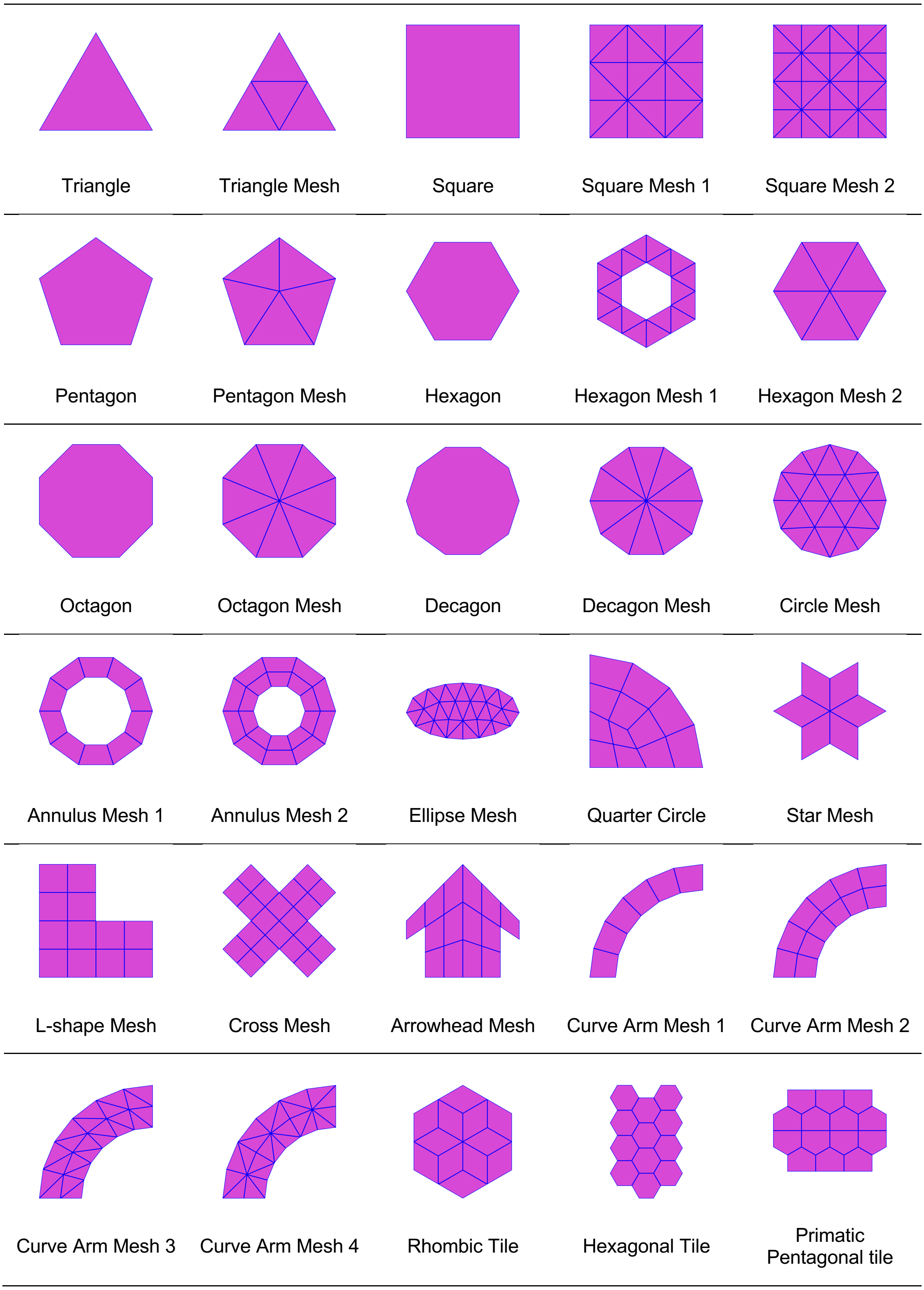

| | | | | |
|---|---|---|---|---|
| 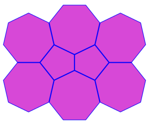 | 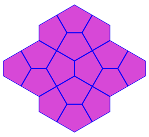 | 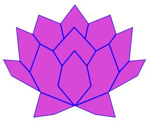 | 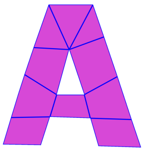 | 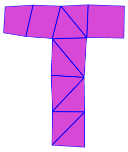 |
| Heptagonal and Pentagonal Tile | Cairo Pentagonal Tile | Lotus Mesh | A-Shape Mesh | T-Shape Mesh |
| 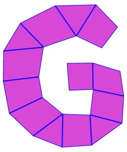 | 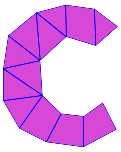 | | | |
| G-Shape Mesh | C-Shape Mesh | | | |

**Supplementary Note 3.** 55 pre-defined 3D target geometries.

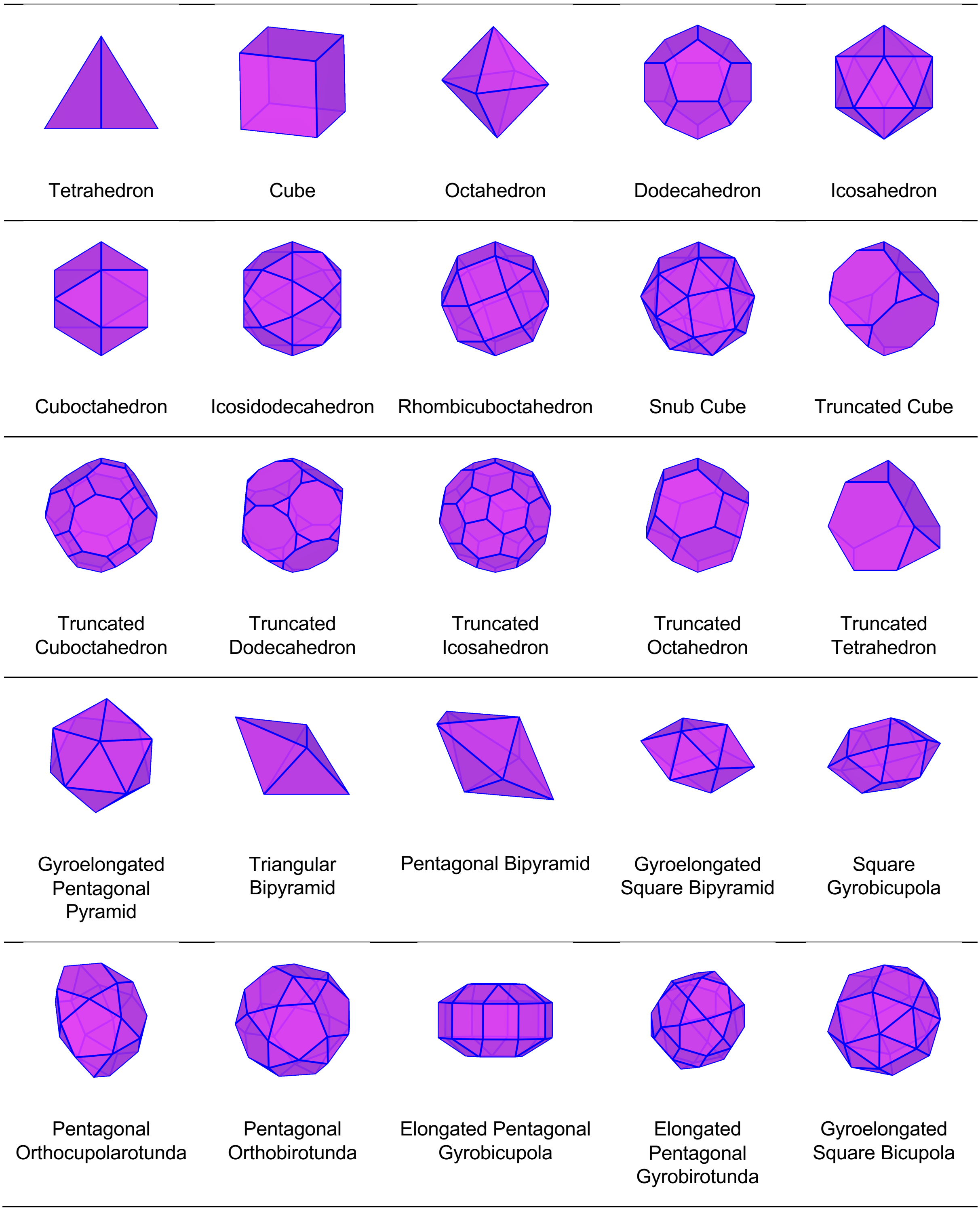

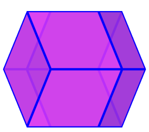
Rhombic Dodecahedron Dodecahedron

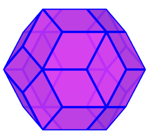
Rhombic Triacontahedron

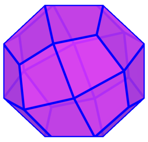
Deltoidal Icositetrahedron

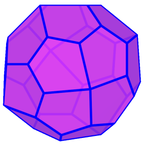
Pentagonal Icositetrahedron

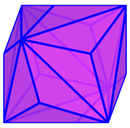
Triakis Octahedron

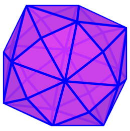
Disdyakis Dodecahedron

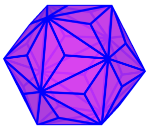
Triakis Icosahedron

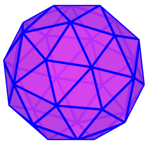
Pentakis Dodecahedron

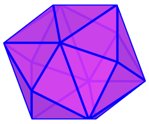
Tetrakis Hexahedron

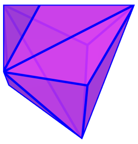
Triakis Tetrahedron

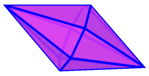
Heptagonal Bipyramid

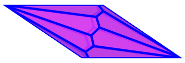
Enneagonal Trapezohedron

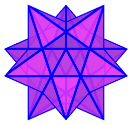
Small Stell Dodecahedron

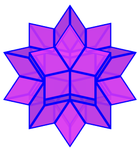
Rhombic Hexecontahedron

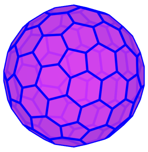
Goldberg Polyhedron dk5dgD

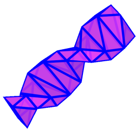
Double Helix

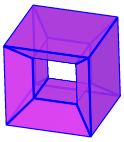
Nested Cube

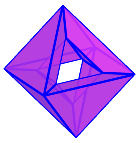
Nested Octahedron

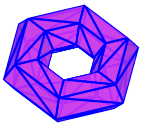
Torus

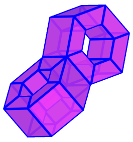
Double Torus

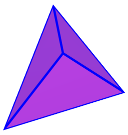
Asymmetric Tetrahedron

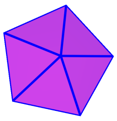
Pentagonal Pyramid

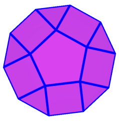
Pentagonal Cupola

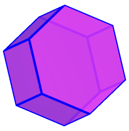
Hexagonal Prism

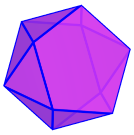
Pentagonal Antiprism

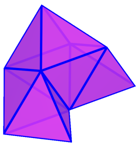
Chiral Object

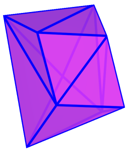
Twisted Triangular Prism

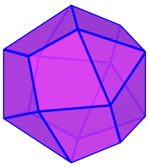
Biscribed Propello Tetrahedron

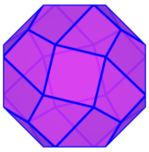
Biscribed Propello Cube

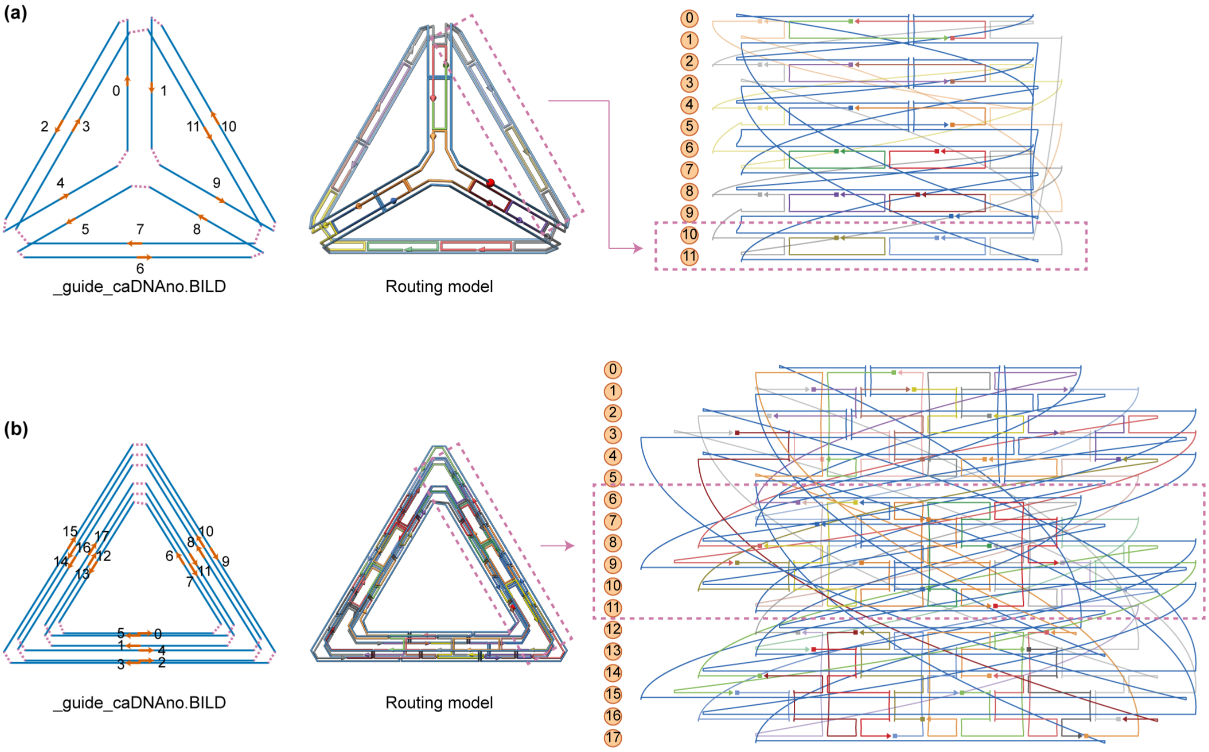


**Figure S1.** Exported SVG schematic of the DNA origami objects. (**a**) DX tetrahedron DNA origami with 63-bp edge-length (**b**) 6HB triangle DNA origami with 63-bp edge-length. Line numbers in the guide model (right) corresponds to row numbers in caDNAno and arrows (orange) indicate the direction of the scaffold. caDNAno representation with scaffold (blue) and staple(multi-color) strands.

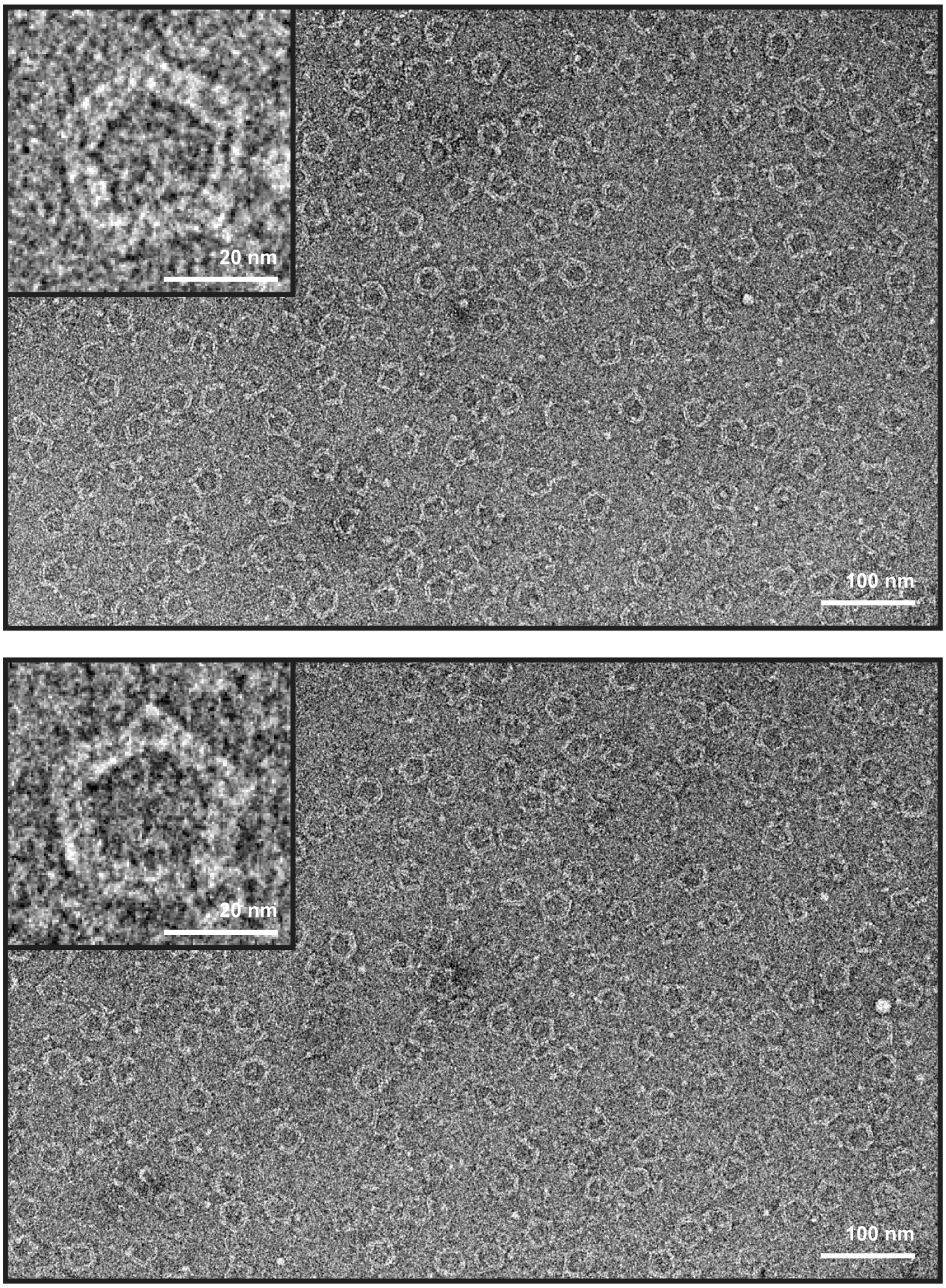


**Figure S2.** TEM imaging of 6HB-based pentagonal DNA origami of 42-bp edge-length without internal mesh.

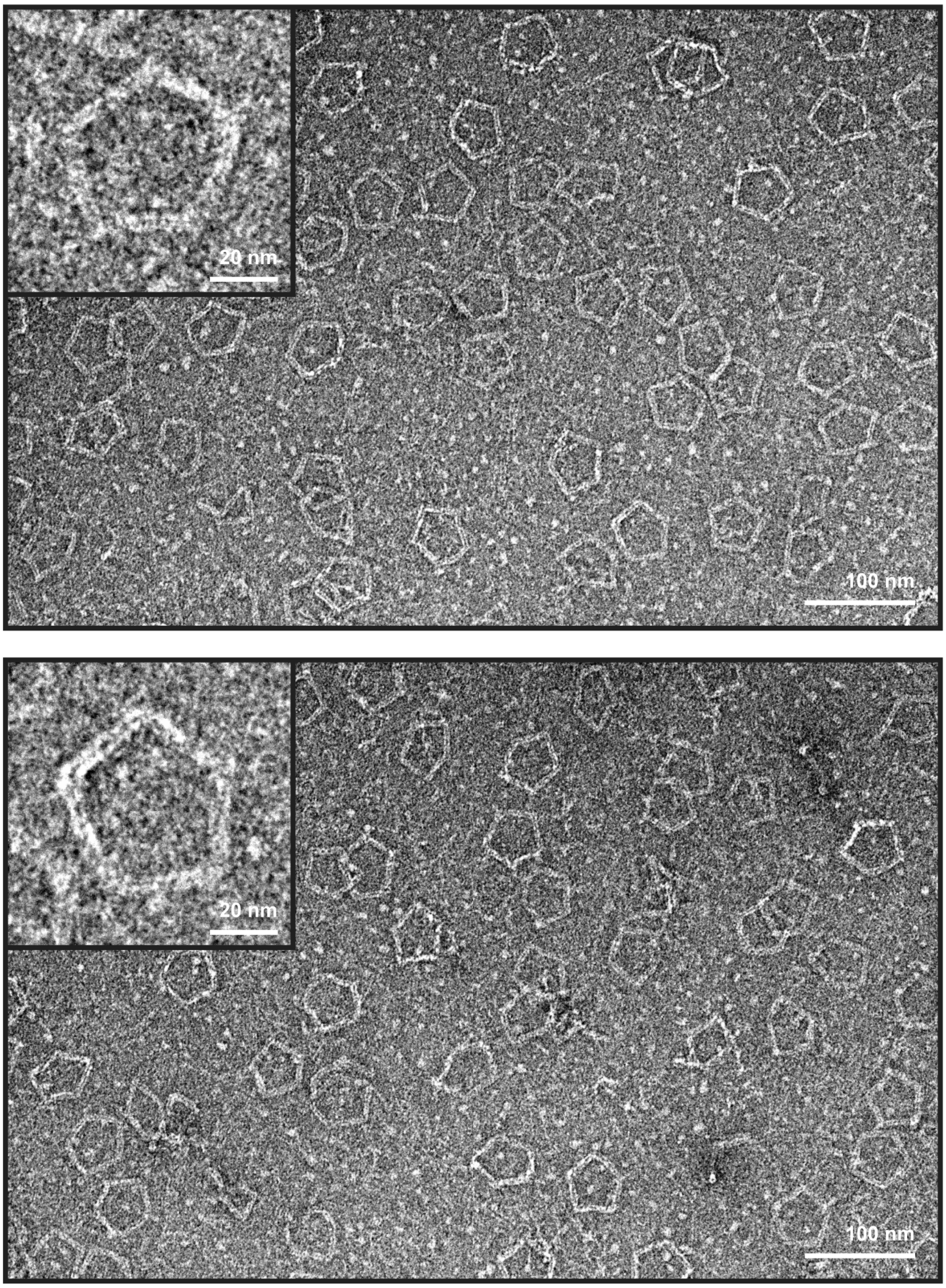


**Figure S3.** TEM imaging of 6HB-based pentagonal DNA origami of 84-bp edge-length without internal mesh.

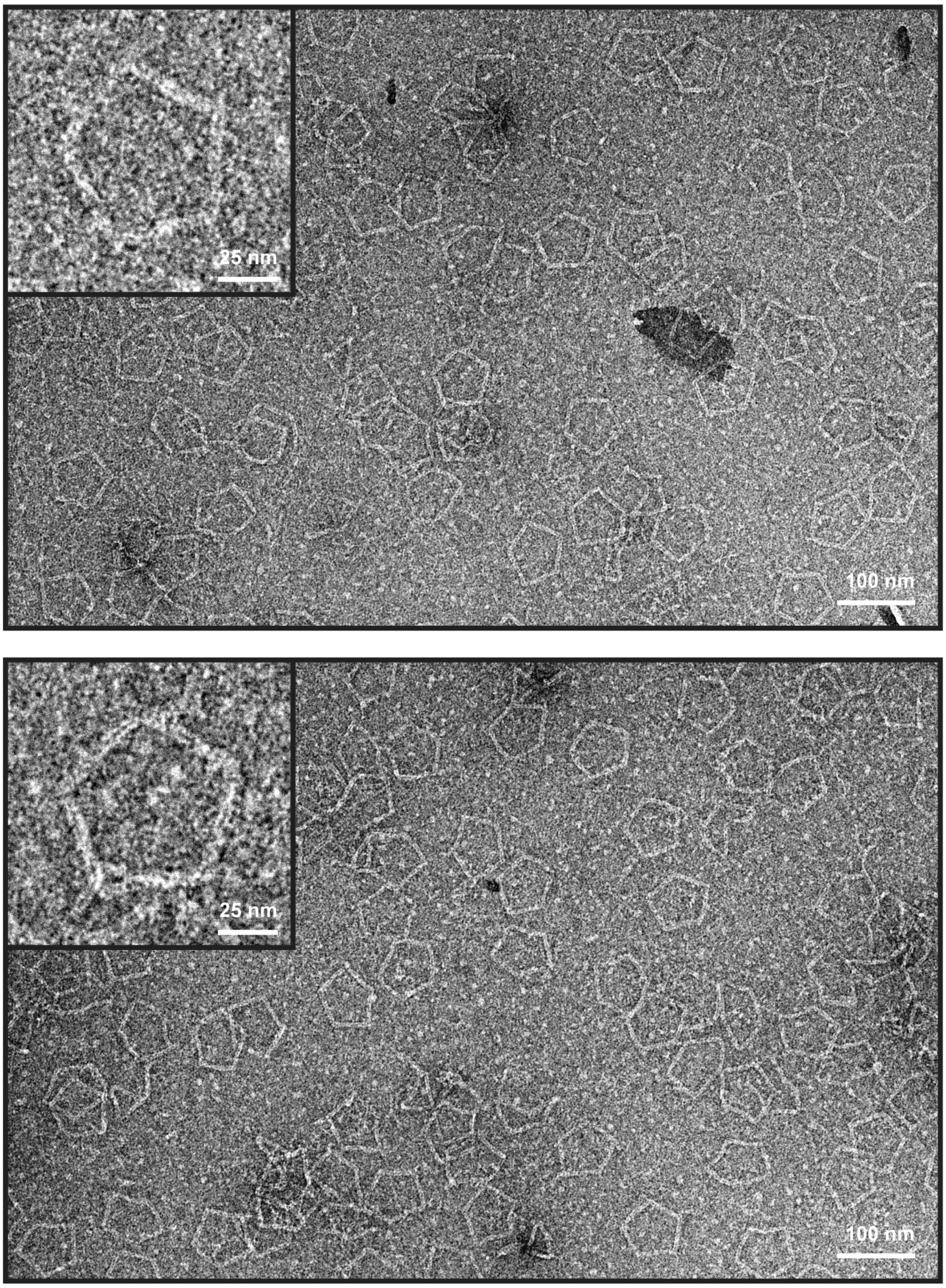


**Figure S4.** TEM imaging of 6HB-based pentagonal DNA origami of 126-bp edge-length without internal mesh.

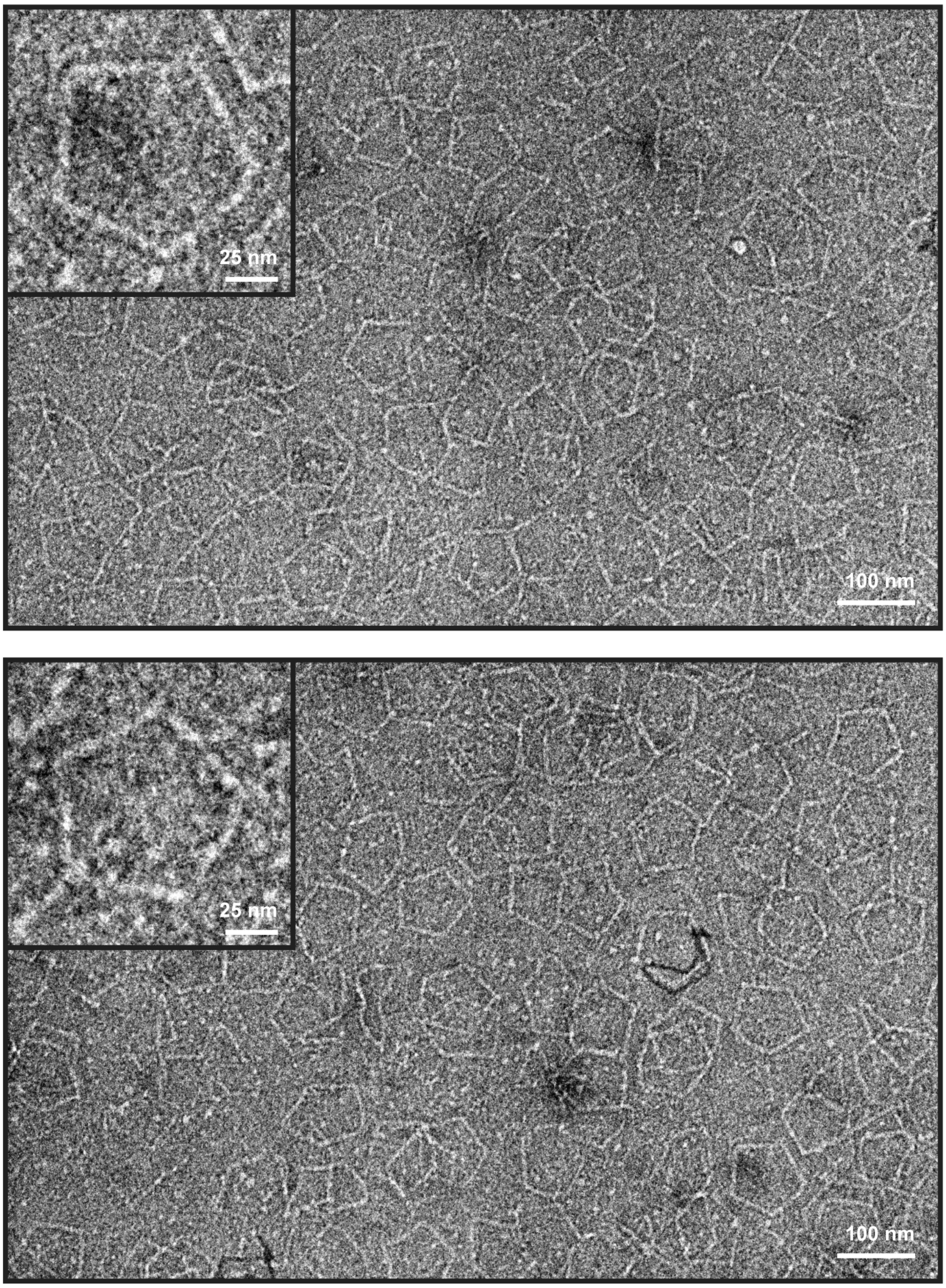


**Figure S5.** TEM imaging of 6HB-based pentagonal DNA origami of 168-bp edge-length without internal mesh.

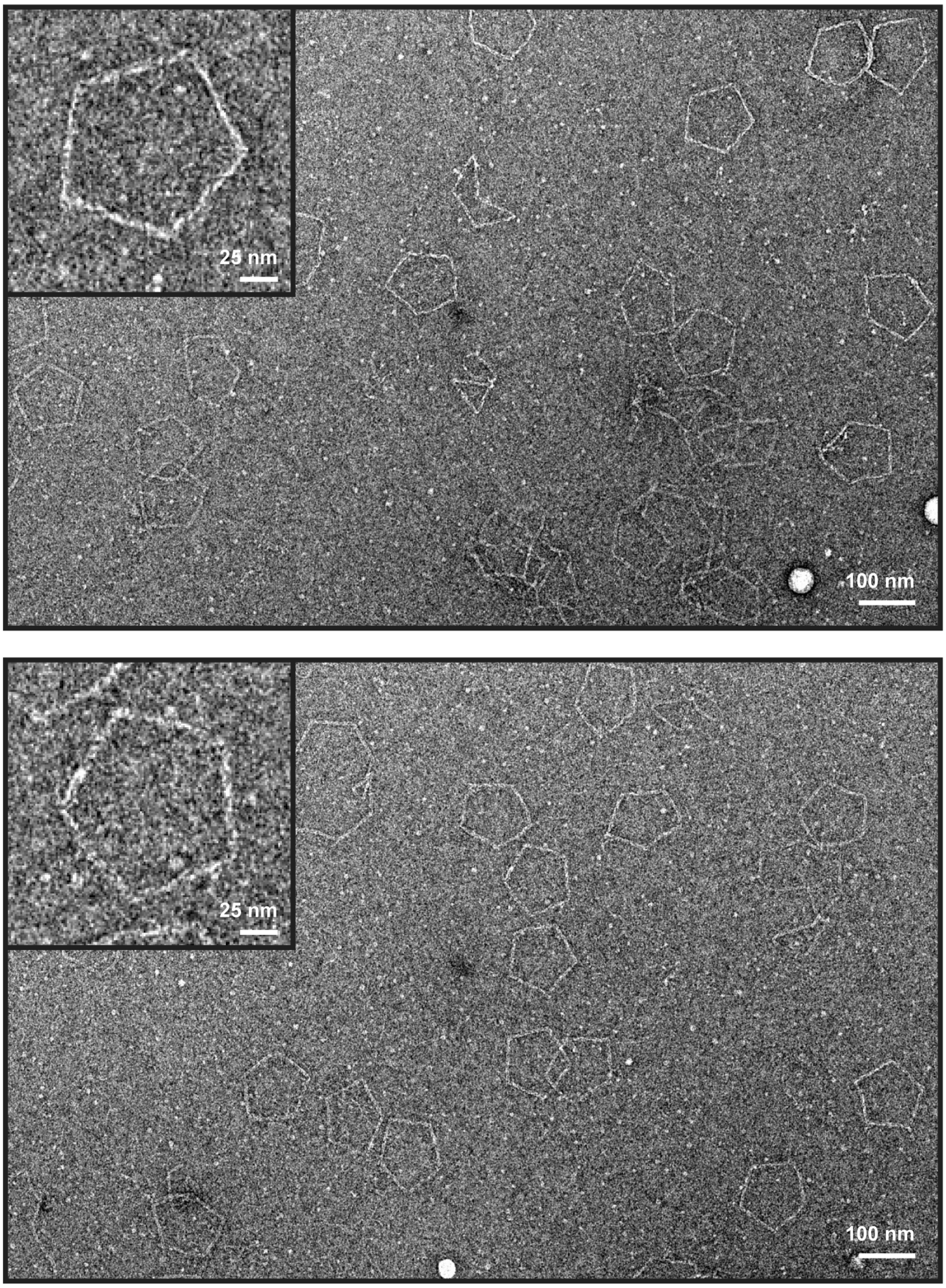


**Figure S6.** TEM imaging of 6HB-based pentagonal DNA origami of 210-bp edge-length without internal mesh.

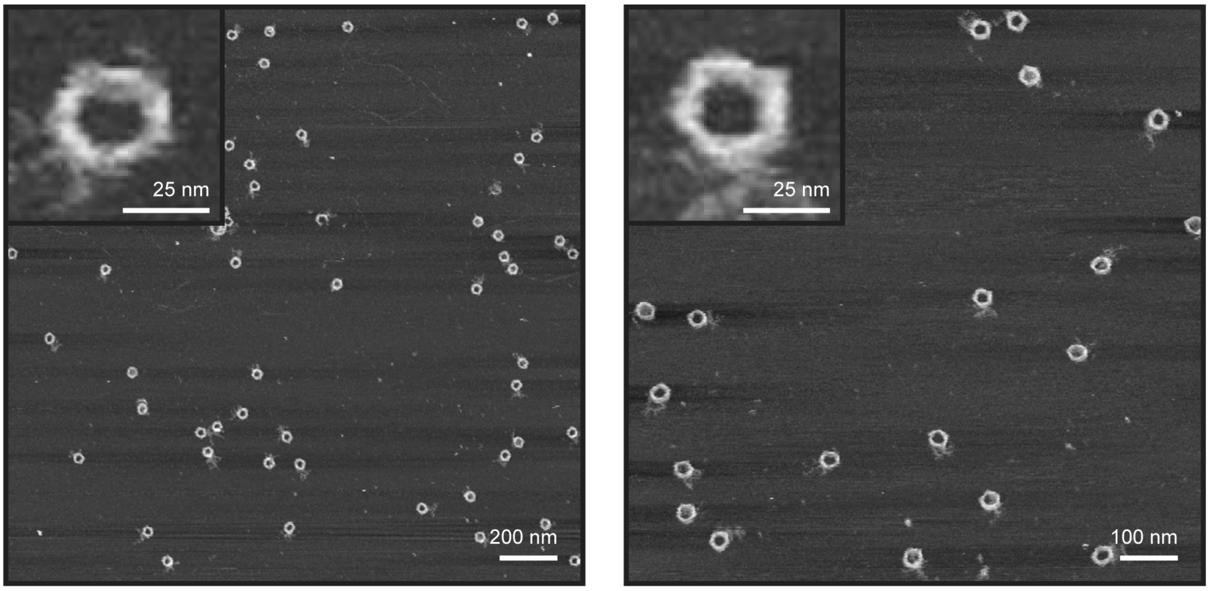


**Figure S7.** AFM imaging of 6HB-based pentagonal DNA origami of 42-bp edge-length without internal mesh.

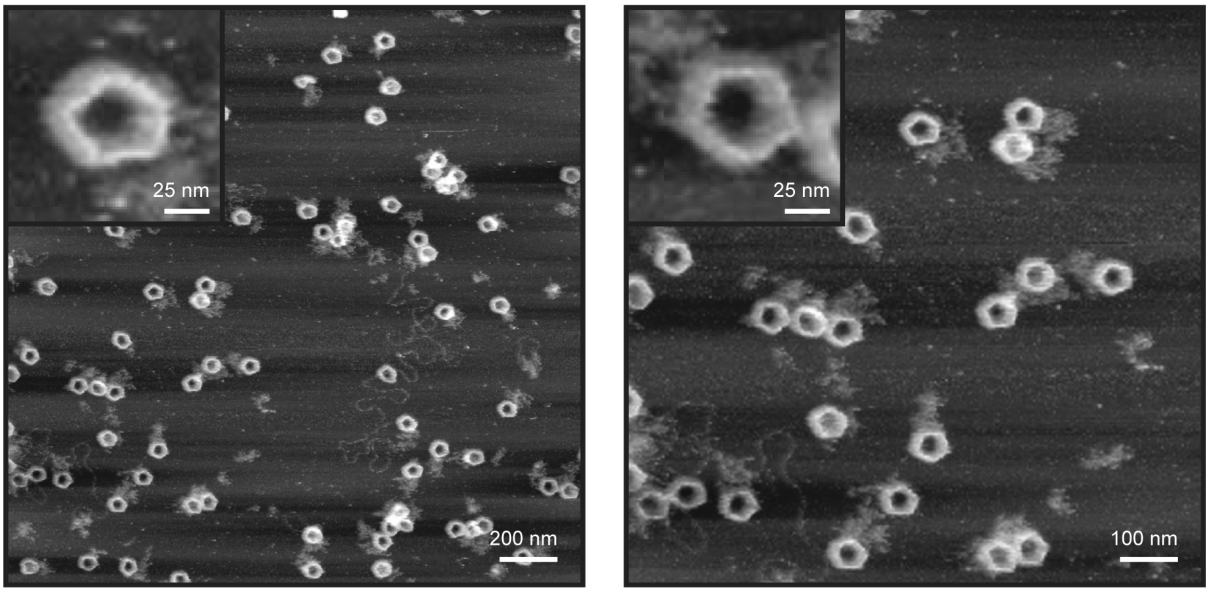


**Figure S8.** AFM imaging of 6HB-based pentagonal DNA origami of 84-bp edge-length without internal mesh.

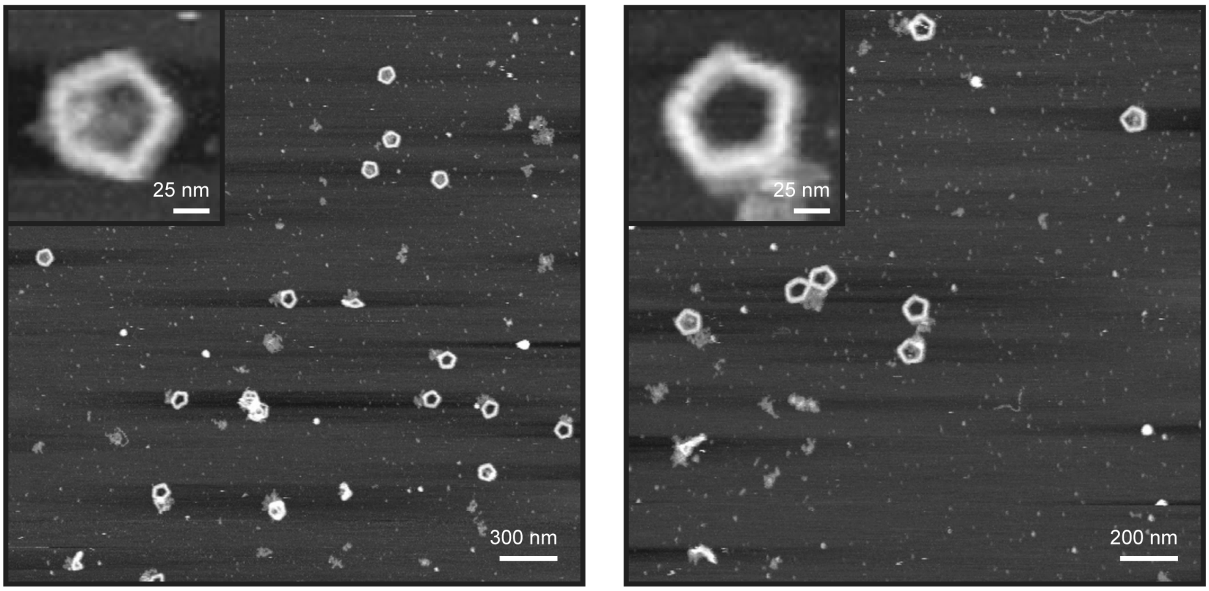


**Figure S9.** AFM imaging of 6HB-based pentagonal DNA origami of 126-bp edge-length without internal mesh.

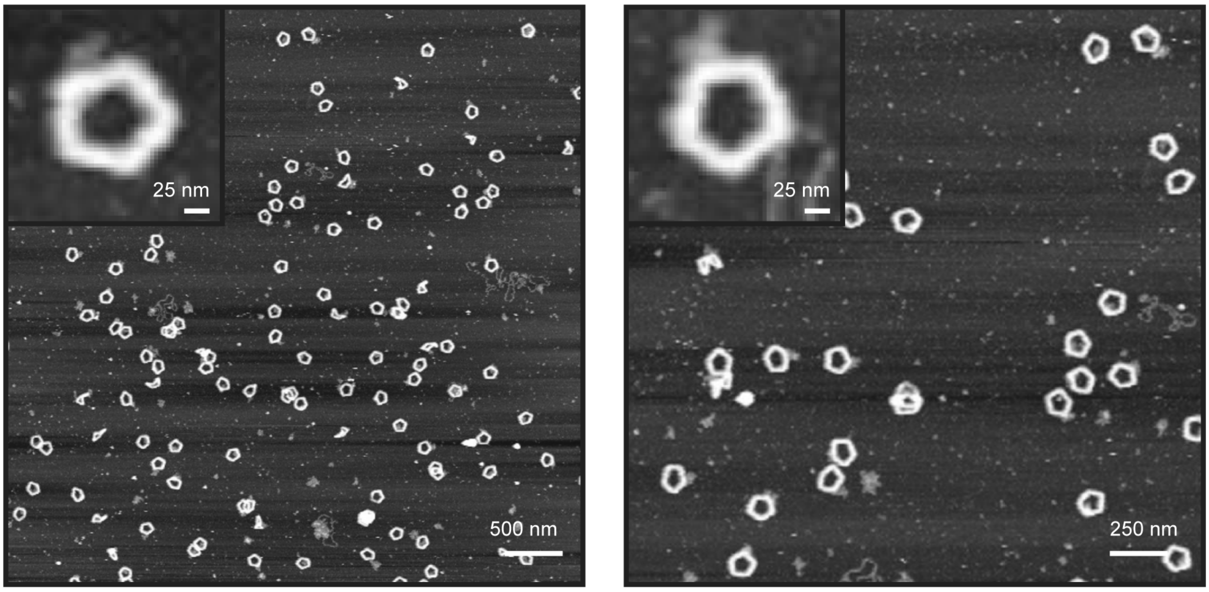


**Figure S10.** AFM imaging of 6HB-based pentagonal DNA origami of 168-bp edge-length without internal mesh.

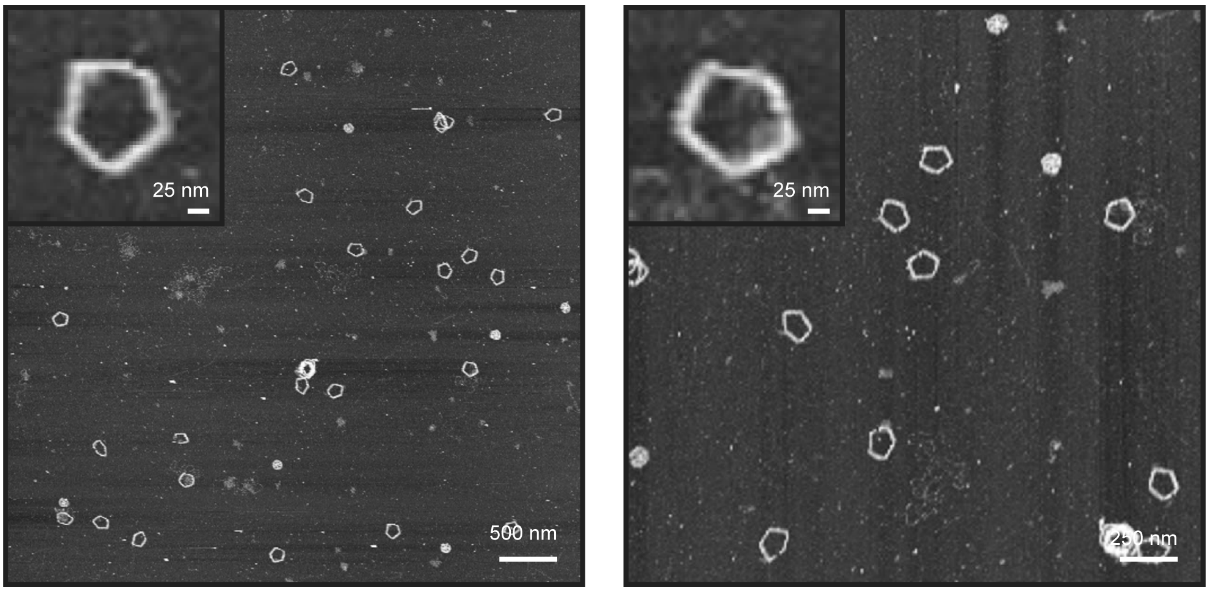


**Figure S11.** AFM imaging of 6HB-based pentagonal DNA origami of 210-bp edge-length without internal mesh.

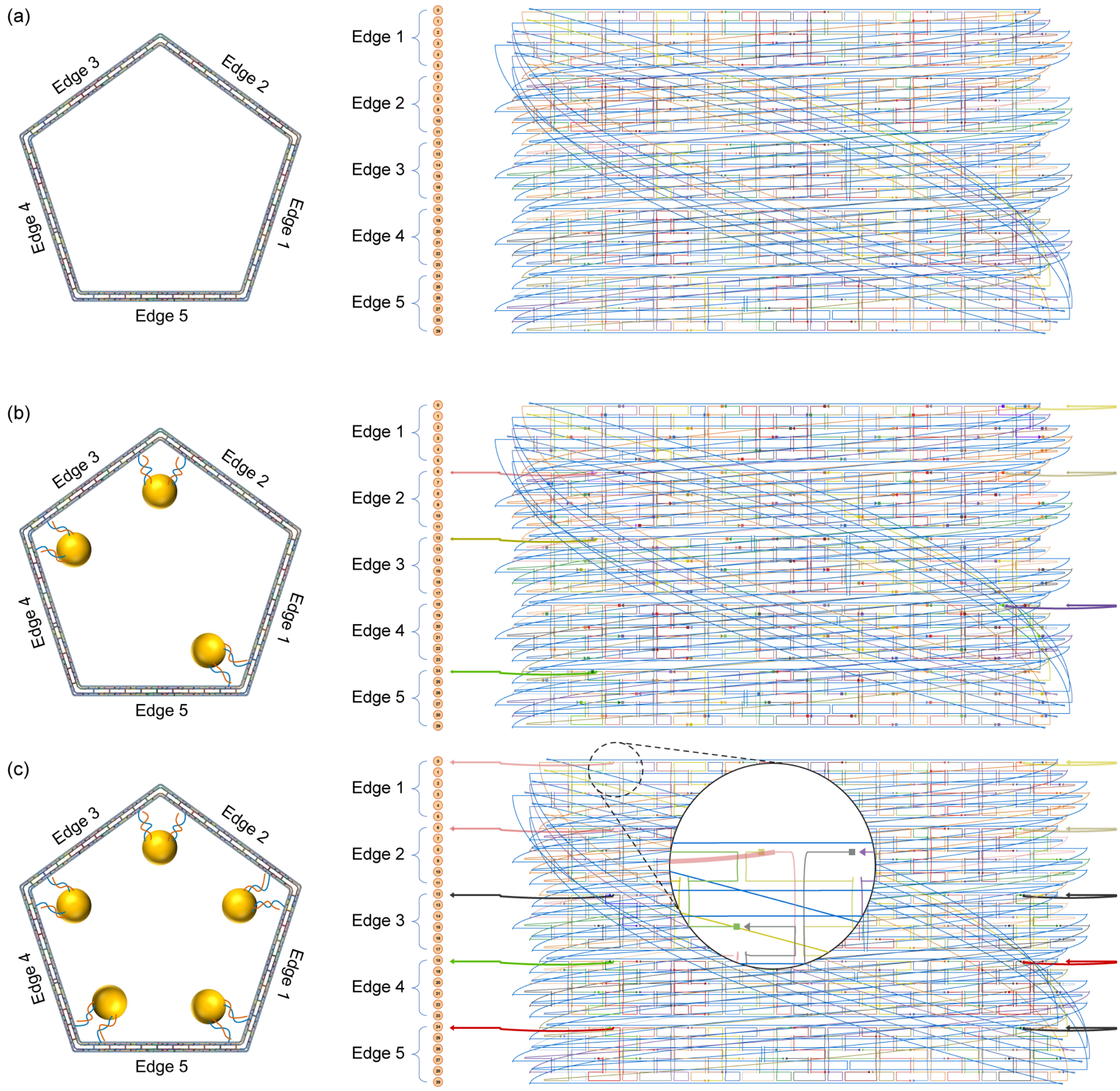


**Figure S12.** Editing origami objects for attaching nanogolds. (**a**) 6HB pentagonal DNA origami with 210-bp edge-length (**b**) 210-bp pentagonal DNA origami with extended staple strands for (**b**) three and (c) five nanogolds. caDNAno representation with scaffold (blue) and staple(multi-color) strands.

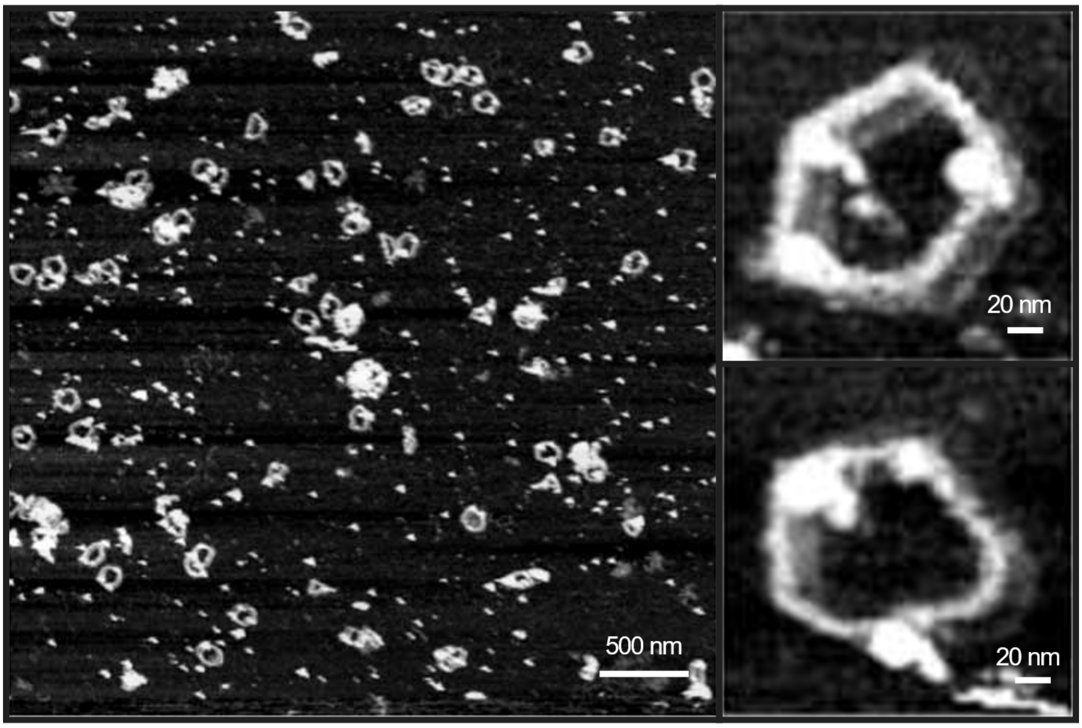


**Figure S13.** AFM imaging of 6HB-based pentagonal DNA origami of 210-bp edge-length three nanogolds.

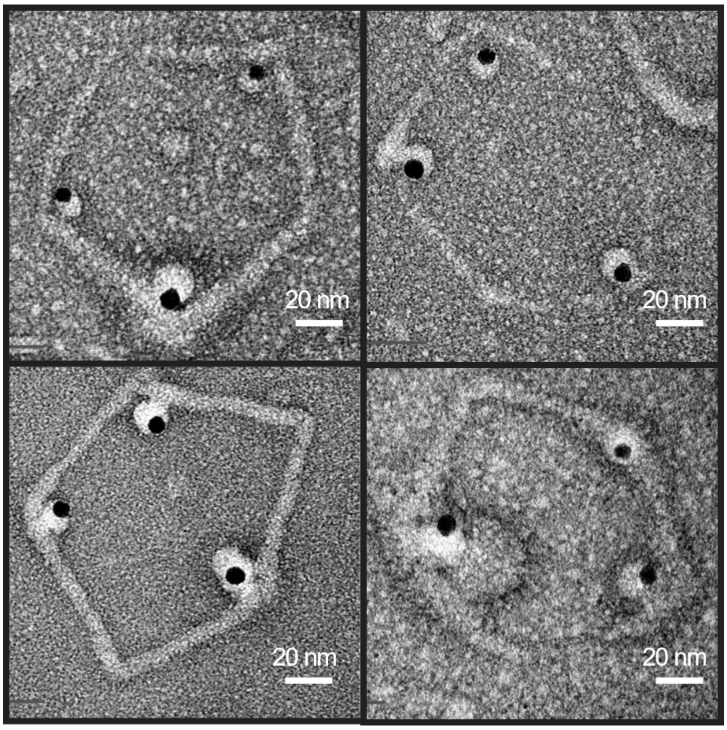


**Figure S14.** TEM imaging of 6HB-based pentagonal DNA origami of 210-bp edge-length with three nanogolds.

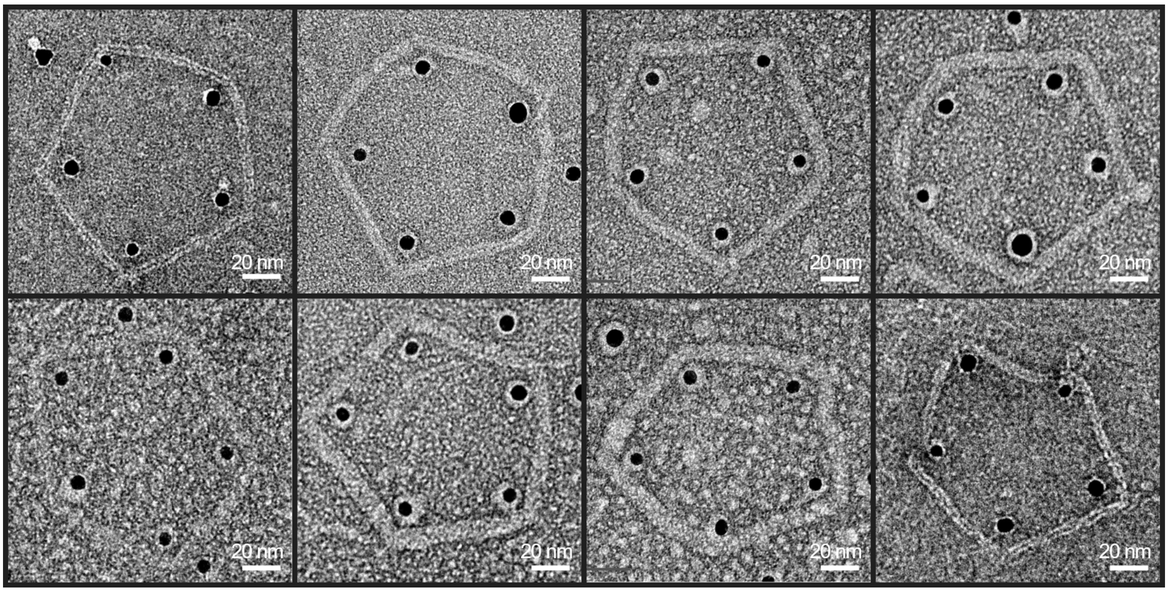


**Figure S15.** TEM imaging of 6HB-based pentagonal DNA origami of 210-bp edge-length with five nanogolds.

**Table S1**. Design parameters for pentagonal DNA origami objects.

| Edge length | Scaffold | | | Staples | | |
|---|---|---|---|---|---|---|
| | Required length | # of double crossovers | # of unpaired nucleotides | # of staples | # of double crossovers | # of unpaired nucleotides |
| **42-bp** | 1,648-nt (#1) | 5 | 133 | 38 | 52 | 60 |
| **84-bp** | 2,908-nt (#2) | 5 | 133 | 68 | 110 | 60 |
| **126-bp** | 4,168-nt (#2) | 5 | 133 | 96 | 170 | 60 |
| **168-bp** | 5,428-nt (#2) | 5 | 133 | 125 | 230 | 60 |
| **210-bp** | 6,688-nt (#2) | 5 | 133 | 157 | 290 | 60 |

### indicates the type of scaffolds in **Table S2**.

**Table S2**. Sequences for the 2,775-nt (#1) and 7,249-nt (#2) length scaffolds used.

| No | Scaffold sequence |
|---|---|
| **#1** | GAGCGCAACGCAATTAATGTGCGCCCTGTAGCGGCGCATTAAGCGCGGCGGGTGTGGTGGTTACGCGCAGCGTGACCGCTAC<br>ACTTGCCAGCGCCCTAGCGCCCGCTCCTTTCGCTTTCTTCCCTTCCTTTCTCGCCACGTTCGCCGGCTTTCCCCGTCAAGCT<br>CTAAATCGGGGGCTCCCTTTAGGGTTCCGATTTAGTGCTTTACGGCACCTCGACCCCAAAAAACTTGATTAGGGTGATGGTT<br>CACGTAGTGGGCCATCGCCCTGATAGACGGTTTTTCGCCCTTTGACGTTGGAGTCCACGTTCTTTAATAGTGGACTCTTGTT<br>CCAAACTGGAACAACACTCAACCCTATCTCGGTCTATTCTTTTGATTTATAAGGGATTTTGCCGATTTCGGCCTATTGGTTA<br>AAAAATGAGCTGATTTAACAAAAATTTAACGCGAATTACAACCGGGGTACATATGATTGGGGTCTGACGCTCAGTGGAACGA<br>AAACTCACGTTAAGGGATTTTGGTCATGAGATTATCAAAAAGGATCTTCACCTAGATCCTTTTAAATTAAAAATGAAGTTTT<br>AAATCAATCTAAAGTATATATGAGTAAACTTGGTCTGACAGTTACCAATGCTTAATCAGTGAGGCACCTATCTCAGCGATCT<br>GTCTATTTCGTTCATCCATAGTTGCCTGACTCCCCGTCGTGTAGATAACTACGATACGGGAGGGCTTACCATCTGGCCCCAG<br>TGCTGCAATGATACCGCGAGACCCACGCTCACCGGCTCCAGATTTATCAGCAATAAACCAGCCAGCCGGAAGGGCCGAGCGC<br>AGAAGTGGTCCTGCAACTTTATCCGCCTCCATCCAGTCTATTAATTGTTGCCGGGAAGCTAGAGTAAGTAGTTCGCCAGTTA<br>ATAGTTTGCGCAACGTTGTTGCCATTGCTACAGGCATCGTGGTGTCACGCTCGTCGTTTGGTATGGCTTCATTCAGCTCCGG<br>TTCCCAACGATCAAGGCGAGTTACATGATCCCCCATGTTGTGCAAAAAAGCGGTTAGCTCCTTCGGTCCTCCGATCGTTGTC<br>AGAAGTAAGTTGGCCGCAGTGTTATCACTCATGGTTATGGCAGCACTGCATAATTCTCTTACTGTCATGCCATCCGTAAGAT<br>GCTTTTCTGTGACTGGTGAGTACTCAACCAAGTCATTCTGAGAATAGTGTATGCGGCGACCGAGTTGCTCTTGCCCGGCGTC<br>AATACGGGATAATACCGCGCCACATAGCAGAACTTTAAAAGTGCTCATCATTGGAAAACGTTCTTCGGGGCGAAAACTCTCA<br>AGGATCTTACCGCTGTTGAGATCCAGTTCGATGTAACCCACTCGTGCACCCAACTGATCTTCAGCATCTTTTACTTTCACCA<br>GCGTTTCTGGGTGAGCAAAAACAGGAAGGCAAAATGCCGCAAAAAAGGGAATAAGGGCGACACGGAAATGTTGAATACTCAT<br>ACTCTTCCTTTTTCAATATTATTGAAGCATTTATCAGGGTTATTGTCTCATGAGCGGATACATATTTGAATGTATTTAGAAA<br>AATAAACAAATAGGGGTTCCGCGCACATTTCCCCGAAAAGTGCCACCTGACGTCTAAGAAACCATTATTATCATGACATTAA<br>CCTATAAAAATAGGCGTATCACGAGGCCCTTTCGTCGAATTCGTCGTCGTCCCCTCAAACTCTTGGGTGGAGAGGCTATTCG<br>TTTAAGGTCACATCGCATGTAATTTACTTATTCTCTGTTGTTGAGCCACCCGGGCGCCAGATTTTGTTTAAAGCTTTGTCTC<br>TTAGTTTGTATAGACAGATTCAGAGTGCAAGGTTTCGTTCGCTCGTACCTGGTTTTCCCTGGTTCTTCACAGATAGGATTTG<br>ACTTTCTACAACACTTATGCGGCTTCCTACCCGTTTGAAGGCCGATACAGGTGCTGCGCAAAATGCGGGCGAACATAGAGTA<br>TCAAAACAACGCCTTCTAATCTAGGAATATAGGGAAGATACGTATTTGCTACCATGCTTTCTGGGGTCATTAACGACCAACC<br>TCTTTTCTTTTAAAGTAGGATTGCACAATGAATGAATACACGTGGTCCGATAACTGACCAAGTAACATGGTTATCACTCGAT<br>GTCCGCCAGACGTGTGCAAACCAACCCGGGAGTTACGTCACTAATCCTTCGCTACGTCGTGAAGATATTTACTTGTGAATAT<br>CGAGGGTAATAAGATAATAGACTGTGACTAGTATTGCCAGACTGTCGCTACCTGCAACACATAACTATCCTGAGGTTACTGC<br>ATAGTACTGATTACACCCGAGTCAAAATTTCTAACTTCTAACATGTACCTAGTAACCAGCTCAATAATTATGTCAGAATATA<br>GCTCTGGGAACCCTCGGACAATTATGATACACGGTATTAATATCTTGCTTGCGTTAGCCACTTCTCATCTTTGGATACCGAT |

| | |
|---|---|
| | TCTATTTTGCATAGCAGTTCCTTTTACACATATAAGAATTTCGCCATAGGTATGACCTACCCCAGATCGTCGATTATCTGCT<br>GGAAAATTTATTTAACACTATGTTTCTCTCCAGATGTGAGTATACACGATAAATAATACCTGGGTACCGGTTGGTGTTATTA<br>CCTTGTTTCTAAGTGCTTAATCGGCGCTTAGTGATAAGGTTGTACTAGTCGACGCGTGGCCGCCAATTATTCTTGTCATAAT<br>TTGACTTTGTTCTATATGACTATGATCTCCTGTCATCTCACCTATTGATGCCACCTTTTCAGCCTGCAG |
| **#2** | AATGCTACTACTATTAGTAGAATTGATGCCACCTTTTCAGCTCGCGCCCCAAATGAAAATATAGCTAAACAGGTTATTGACC<br>ATTTGCGAAATGTATCTAATGGTCAAACTAAATCTACTCGTTCGCAGAATTGGGAATCAACTGTTATATGGAATGAAACTTC<br>CAGACACCGTACTTTAGTTGCATATTTAAAACATGTTGAGCTACAGCATTATATTCAGCAATTAAGCTCTAAGCCATCCGCA<br>AAAATGACCTCTTATCAAAAGGAGCAATTAAAGGTACTCTCTAATCCTGACCTGTTGGAGTTTGCTTCCGGTCTGGTTCGCT<br>TTGAAGCTCGAATTAAAACGCGATATTTGAAGTCTTTCGGGCTTCCTCTTAATCTTTTTGATGCAATCCGCTTTGCTTCTGA<br>CTATAATAGTCAGGGTAAAGACCTGATTTTTGATTTATGGTCATTCTCGTTTTCTGAACTGTTTAAAGCATTTGAGGGGGAT<br>TCAATGAATATTTATGACGATTCCGCAGTATTGGACGCTATCCAGTCTAAACATTTTACTATTACCCCCTCTGGCAAAACTT<br>CTTTTGCAAAAGCCTCTCGCTATTTTGGTTTTTATCGTCGTCTGGTAAACGAGGGTTATGATAGTGTTGCTCTTACTATGCC<br>TCGTAATTCCTTTTGGCGTTATGTATCTGCATTAGTTGAATGTGGTATTCCTAAATCTCAACTGATGAATCTTTCTACCTGT<br>AATAATGTTGTTCCGTTAGTTCGTTTTATTAACGTAGATTTTTCTTCCCAACGTCCTGACTGGTATAATGAGCCAGTTCTTA<br>AAATCGCATAAGGTAATTCACAATGATTAAAGTTGAAATTAAACCATCTCAAGCCCAATTTACTACTCGTTCTGGTGTTTCT<br>CGTCAGGGCAAGCCTTATTCACTGAATGAGCAGCTTTGTTACGTTGATTTGGGTAATGAATATCCGGTTCTTGTCAAGATTA<br>CTCTTGATGAAGGTCAGCCAGCCTATGCGCCTGGTCTGTACACCGTTCATCTGTCCTCTTTCAAAGTTGGTCAGTTCGGTTC<br>CCTTATGATTGACCGTCTGCGCCTCGTTCCGGCTAAGTAACATGGAGCAGGTCGCGGATTTCGACACAATTTATCAGGCGAT<br>GATACAAATCTCCGTTGTACTTTGTTTCGCGCTTGGTATAATCGCTGGGGGTCAAAGATGAGTGTTTTAGTGTATTCTTTTG<br>CCTCTTTCGTTTTAGGTTGGTGCCTTCGTAGTGGCATTACGTATTTTACCCGTTTAATGGAAACTTCCTCATGAAAAAGTCT<br>TTAGTCCTCAAAGCCTCTGTAGCCGTTGCTACCCTCGTTCCGATGCTGTCTTTCGCTGCTGAGGGTGACGATCCCGCAAAAG<br>CGGCCTTTAACTCCCTGCAAGCCTCAGCGACCGAATATATCGGTTATGCGTGGGCGATGGTTGTTGTCATTGTCGGCGCAAC<br>TATCGGTATCAAGCTGTTTAAGAAATTCACCTCGAAAGCAAGCTGATAAACCGATACAATTAAAGGCTCCTTTTGGAGCCTT<br>TTTTTTGGAGATTTTCAACGTGAAAAAATTATTATTCGCAATTCCTTTAGTTGTTCCTTTCTATTCTCACTCCGCTGAAACT<br>GTTGAAAGTTGTTTAGCAAAATCCCATACAGAAAATTCATTTACTAACGTCTGGAAAGACGACAAAACTTTAGATCGTTACG<br>CTAACTATGAGGGCTGTCTGTGGAATGCTACAGGCGTTGTAGTTTGTACTGGTGACGAAACTCAGTGTTACGGTACATGGGT<br>TCCTATTGGGCTTGCTATCCCTGAAAATGAGGGTGGTGGCTCTGAGGGTGGCGGTTCTGAGGGTGGCGGTTCTGAGGGTGGC<br>GGTACTAAACCTCCTGAGTACGGTGATACACCTATTCCGGGCTATACTTATATCAACCCTCTCGACGGCACTTATCCGCCTG<br>GTACTGAGCAAAACCCCGCTAATCCTAATCCTTCTCTTGAGGAGTCTCAGCCTCTTAATACTTTCATGTTTCAGAATAATAG<br>GTTCCGAAATAGGCAGGGGGCATTAACTGTTTATACGGGCACTGTTACTCAAGGCACTGACCCCGTTAAAACTTATTACCAG<br>TACACTCCTGTATCATCAAAAGCCATGTATGACGCTTACTGGAACGGTAAATTCAGAGACTGCGCTTTCCATTCTGGCTTTA<br>ATGAGGATTTATTTGTTTGTGAATATCAAGGCCAATCGTCTGACCTGCCTCAACCTCCTGTCAATGCTGGCGGCGGCTCTGG<br>TGGTGGTTCTGGTGGCGGCTCTGAGGGTGGTGGCTCTGAGGGTGGCGGTTCTGAGGGTGGCGGCTCTGAGGGAGGCGGTTCC<br>GGTGGTGGCTCTGGTTCCGGTGATTTTGATTATGAAAAGATGGCAAACGCTAATAAGGGGGCTATGACCGAAAATGCCGATG<br>AAAACGCGCTACAGTCTGACGCTAAAGGCAAACTTGATTCTGTCGCTACTGATTACGGTGCTGCTATCGATGGTTTCATTGG<br>TGACGTTTCCGGCCTTGCTAATGGTAATGGTGCTACTGGTGATTTTGCTGGCTCTAATTCCCAAATGGCTCAAGTCGGTGAC<br>GGTGATAATTCACCTTTAATGAATAATTTCCGTCAATATTTACCTTCCCTCCCTCAATCGGTTGAATGTCGCCCTTTTGTCT<br>TTGGCGCTGGTAAACCATATGAATTTTCTATTGATTGTGACAAAATAAACTTATTCCGTGGTGTCTTTGCGTTTCTTTTATA<br>TGTTGCCACCTTTATGTATGTATTTTCTACGTTTGCTAACATACTGCGTAATAAGGAGTCTTAATCATGCCAGTTCTTTTGG<br>GTATTCCGTTATTATTGCGTTTCCTCGGTTTCCTTCTGGTAACTTTGTTCGGCTATCTGCTTACTTTTCTTAAAAAGGGCTT<br>CGGTAAGATAGCTATTGCTATTTCATTGTTTCTTGCTCTTATTATTGGGCTTAACTCAATTCTTGTGGGTTATCTCTCTGAT<br>ATTAGCGCTCAATTACCCTCTGACTTTGTTCAGGGTGTTCAGTTAATTCTCCCGTCTAATGCGCTTCCCTGTTTTTATGTTA<br>TTCTCTCTGTAAAGGCTGCTATTTTCATTTTTGACGTTAAACAAAAAATCGTTTCTTATTTGGATTGGGATAAATAATATGG<br>CTGTTTATTTTGTAACTGGCAAATTAGGCTCTGGAAAGACGCTCGTTAGCGTTGGTAAGATTCAGGATAAAATTGTAGCTGG<br>GTGCAAAATAGCAACTAATCTTGATTTAAGGCTTCAAAACCTCCCGCAAGTCGGGAGGTTCGCTAAAACGCCTCGCGTTCTT<br>AGAATACCGGATAAGCCTTCTATATCTGATTTGCTTGCTATTGGGCGCGGTAATGATTCCTACGATGAAAATAAAAACGGCT<br>TGCTTGTTCTCGATGAGTGCGGTACTTGGTTTAATACCCGTTCTTGGAATGATAAGGAAAGACAGCCGATTATTGATTGGTT<br>TCTACATGCTCGTAAATTAGGATGGGATATTATTTTTCTTGTTCAGGACTTATCTATTGTTGATAAACAGGCGCGTTCTGCA<br>TTAGCTGAACATGTTGTTTATTGTCGTCGTCTGGACAGAATTACTTTACCTTTTGTCGGTACTTTATATTCTCTTATTACTG<br>GCTCGAAAATGCCTCTGCCTAAATTACATGTTGGCGTTGTTAAATATGGCGATTCTCAATTAAGCCCTACTGTTGAGCGTTG<br>GCTTTATACTGGTAAGAATTTGTATAACGCATATGATACTAAACAGGCTTTTTCTAGTAATTATGATTCCGGTGTTTATTCT<br>TATTTAACGCCTTATTTATCACACGGTCGGTATTTCAAACCATTAAATTTAGGTCAGAAGATGAAATTAACTAAAATATATT<br>TGAAAAAGTTTTCTCGCGTTCTTTGTCTTGCGATTGGATTTGCATCAGCATTTACATATAGTTATATAACCCAACCTAAGCC<br>GGAGGTTAAAAAGGTAGTCTCTCAGACCTATGATTTTGATAAATTCACTATTGACTCTTCTCAGCGTCTTAATCTAAGCTAT<br>CGCTATGTTTTCAAGGATTCTAAGGGAAAATTAATTAATAGCGACGATTTACAGAAGCAAGGTTATTCACTCACATATATTG<br>ATTTATGTACTGTTTCCATTAAAAAAGGTAATTCAAATGAAATTGTTAAATGTAATTAATTTTGTTTTCTTGATGTTTGTTT |

| | |
|---|---|
| | CATCATCTTCTTTTGCTCAGGTAATTGAAATGAATAATTCGCCTCTGCGCGATTTTGTAACTTGGTATTCAAAGCAATCAGG<br>CGAATCCGTTATTGTTTCTCCCGATGTAAAAGGTACTGTTACTGTATATTCATCTGACGTTAAACCTGAAAATCTACGCAAT<br>TTCTTTATTTCTGTTTTACGTGCAAATAATTTTGATATGGTAGGTTCTAACCCTTCCATTATTCAGAAGTATAATCCAAACA<br>ATCAGGATTATATTGATGAATTGCCATCATCTGATAATCAGGAATATGATGATAATTCCGCTCCTTCTGGTGGTTTCTTTGT<br>TCCGCAAAATGATAATGTTACTCAAACTTTTAAAATTAATAACGTTCGGGCAAAGGATTTAATACGAGTTGTCGAATTGTTT<br>GTAAAGTCTAATACTTCTAAATCCTCAAATGTATTATCTATTGACGGCTCTAATCTATTAGTTGTTAGTGCTCCTAAAGATA<br>TTTTAGATAACCTTCCTCAATTCCTTTCAACTGTTGATTTGCCAACTGACCAGATATTGATTGAGGGTTTGATATTTGAGGT<br>TCAGCAAGGTGATGCTTTAGATTTTTCATTTGCTGCTGGCTCTCAGCGTGGCACTGTTGCAGGCGGTGTTAATACTGACCGC<br>CTCACCTCTGTTTTATCTTCTGCTGGTGGTTCGTTCGGTATTTTTAATGGCGATGTTTTAGGGCTATCAGTTCGCGCATTAA<br>AGACTAATAGCCATTCAAAAATATTGTCTGTGCCACGTATTCTTACGCTTTCAGGTCAGAAGGGTTCTATCTCTGTTGGCCA<br>GAATGTCCCTTTTATTACTGGTCGTGTGACTGGTGAATCTGCCAATGTAAATAATCCATTTCAGACGATTGAGCGTCAAAAT<br>GTAGGTATTTCCATGAGCGTTTTTCCTGTTGCAATGGCTGGCGGTAATATTGTTCTGGATATTACCAGCAAGGCCGATAGTT<br>TGAGTTCTTCTACTCAGGCAAGTGATGTTATTACTAATCAAAGAAGTATTGCTACAACGGTTAATTTGCGTGATGGACAGAC<br>TCTTTTACTCGGTGGCCTCACTGATTATAAAAACACTTCTCAGGATTCTGGCGTACCGTTCCTGTCTAAAATCCCTTTAATC<br>GGCCTCCTGTTTAGCTCCCGCTCTGATTCTAACGAGGAAAGCACGTTATACGTGCTCGTCAAAGCAACCATAGTACGCGCCC<br>TGTAGCGGCGCATTAAGCGCGGCGGGTGTGGTGGTTACGCGCAGCGTGACCGCTACACTTGCCAGCGCCCTAGCGCCCGCTC<br>CTTTCGCTTTCTTCCCTTCCTTTCTCGCCACGTTCGCCGGCTTTCCCCGTCAAGCTCTAAATCGGGGGCTCCCTTTAGGGTT<br>CCGATTTAGTGCTTTACGGCACCTCGACCCCAAAAAACTTGATTTGGGTGATGGTTCACGTAGTGGGCCATCGCCCTGATAG<br>ACGGTTTTTCGCCCTTTGACGTTGGAGTCCACGTTCTTTAATAGTGGACTCTTGTTCCAAACTGGAACAACACTCAACCCTA<br>TCTCGGGCTATTCTTTTGATTTATAAGGGATTTTGCCGATTTCGGAACCACCATCAAACAGGATTTTCGCCTGCTGGGGCAA<br>ACCAGCGTGGACCGCTTGCTGCAACTCTCTCAGGGCCAGGCGGTGAAGGGCAATCAGCTGTTGCCCGTCTCACTGGTGAAAA<br>GAAAAACCACCCTGGCGCCCAATACGCAAACCGCCTCTCCCCGCGCGTTGGCCGATTCATTAATGCAGCTGGCACGACAGGT<br>TTCCCGACTGGAAAGCGGGCAGTGAGCGCAACGCAATTAATGTGAGTTAGCTCACTCATTAGGCACCCCAGGCTTTACACTT<br>TATGCTTCCGGCTCGTATGTTGTGTGGAATTGTGAGCGGATAACAATTTCACACAGGAAACAGCTATGACCATGATTACGAA<br>TTCGAGCTCGGTACCCGGGGATCCTCTAGAGTCGACCTGCAGGCATGCAAGCTTGGCACTGGCCGTCGTTTTACAACGTCGT<br>GACTGGGAAAACCCTGGCGTTACCCAACTTAATCGCCTTGCAGCACATCCCCCTTTCGCCAGCTGGCGTAATAGCGAAGAGG<br>CCCGCACCGATCGCCCTTCCCAACAGTTGCGCAGCCTGAATGGCGAATGGCGCTTTGCCTGGTTTCCGGCACCAGAAGCGGT<br>GCCGGAAAGCTGGCTGGAGTGCGATCTTCCTGAGGCCGATACTGTCGTCGTCCCCTCAAACTGGCAGATGCACGGTTACGAT<br>GCGCCCATCTACACCAACGTGACCTATCCCATTACGGTCAATCCGCCGTTTGTTCCCACGGAGAATCCGACGGGTTGTTACT<br>CGCTCACATTTAATGTTGATGAAAGCTGGCTACAGGAAGGCCAGACGCGAATTATTTTTGATGGCGTTCCTATTGGTTAAAA<br>AATGAGCTGATTTAACAAAAATTTAATGCGAATTTTAACAAAATATTAACGTTTACAATTTAAATATTTGCTTATACAATCT<br>TCCTGTTTTTGGGGCTTTTCTGATTATCAACCGGGGTACATATGATTGACATGCTAGTTTTACGATTACCGTTCATCGATTC<br>TCTTGTTTGCTCCAGACTCTCAGGCAATGACCTGATAGCCTTTGTAGATCTCTCAAAAATAGCTACCCTCTCCGGCATTAAT<br>TTATCAGCTAGAACGGTTGAATATCATATTGATGGTGATTTGACTGTCTCCGGCCTTTCTCACCCTTTTGAATCTTTACCTA<br>CACATTACTCAGGCATTGCATTTAAAATATATGAGGGTTCTAAAAATTTTTATCCTTGCGTTGAAATAAAGGCTTCTCCCGC<br>AAAAGTATTACAGGGTCATAATGTTTTTGGTACAACCGATTTAGCTTTATGCTCTGAGGCTTTATTGCTTAATTTTGCTAAT<br>TCTTTGCCTTGCCTGTATGATTTATTGGATGTT |

**Table S3**. Staple sequence for the pentagonal DNA origami of 42-bp edge-length. The sequence is represented by colors; unpaired nucleotides with blue, crossovers with orange, and the 14-nt seed dsDNA domain with green.

| Staple ID | Length (bp) | Staple sequences |
|---|---|---|
| 1 | 58 | CCCTAATCGATGGCCTGCCATAACCATGAAGAACGTACATTCAAATATGTATTCATGA |
| 2 | 56 | TACTCTTTTCCCGGCAACAATTTGCGCAAATAACTGGCGATTTTAAATTTTAAATC |
| 3 | 55 | TTTTCGCAGTAAGAGAATTAGCACTTTCTATTTGTTTATTTGAAATGTACCATCA |
| 4 | 53 | ACTTTTCGGGTTCTAAATTTTCCAATGATGATGCAGTGCCACTACGTCGTGAG |
| 5 | 50 | GGTTGTATCTAGGTTTTACTCATATATAGTAGTTACATACACTAACTACT |
| 6 | 46 | CATGATTCCACTTGTCAGACCCCAATAATCTCATTTTGACCAAAAT |
| 7 | 46 | GGTTGAGTTAGAGCCGCCGCTACAGGGCCGTGTCGAACGCTGGTGA |
| 8 | 45 | ATTTCGCACATTTCGTTGCGCTCTTTATAGGTAATATTAGTGGGT |
| 9 | 45 | GTATCCTTTAGTTTTAAAACTTGGCGCTAGGATTGCAGCTGGATG |
| 10 | 43 | GGATCCCAGTTTTAAGAGTCCACTACCCTAAAGGTTTGAGCCC |
| 11 | 43 | CGAGCAAAATCTTAAATCAAAAGACGGCGAACGTTTTGGCGAG |
| 12 | 43 | AGCTCCGCATTTTTATTTATTTAAAAGGAATTCGCGCTCAGAA |
| 13 | 43 | CTTACGGGCGATTGTCTATCAGGGCAAGTTTTTTTTTGGGGTC |
| 14 | 43 | CTTTTTGATCATATACTCACCAGTCACAACGCCGGCAACTATG |
| 15 | 43 | AACTGTCAAGTCAGGGCAAGAGCAACTCGGTTGAGTGTACCCC |
| 16 | 42 | GGAGCTGGGTGCCTTCCTGTTTTTGCTCGATCGTTGAGATAG |
| 17 | 42 | GAGGCGGCCTGTAGAACCAATAGGCCGAAAGGGAACAAGTGT |
| 18 | 42 | CGGTATCGCGCTGGGAAAGCGAAAGGAGATTTTTTCAATGGC |
| 19 | 42 | TGACTTGGTCGCCGTCTACACGACGGGGGACCAAGGAAGATC |
| 20 | 42 | GCTTCAATTAATGTAGCACTAAATCGGAATTAAAGCGAAGGA |
| 21 | 42 | TACATCGGGAGGACAACGTGGACTCCAAGCCGTAACATGATA |
| 22 | 42 | TCCCTTTCTTAATGTTGACGGGGAAAGCATAGACCGGGAACC |
| 23 | 41 | TTCTGCGCTCTTTTTCCGGCTGGCTAATGAAGCCTAAACGA |
| 24 | 41 | TATGTGGCGCTTTTATCCCGTATTGGAAAAGCATTGGATGG |
| 25 | 41 | GTAAGATCCTTTTGTTTTCGCCCCGAGTGATAACTGGCCAA |
| 26 | 41 | AAGTAAAAGATTTAAGATCAGTTGGGCTTTTTTGTCATGGG |
| 27 | 41 | CCCTTAAGAGCGCGGAACAGGTGCCTCACTTAAGCATTGGT |
| 28 | 40 | ACCCGCCGCGTTTGCGGCATTTTTTATTGCTGATCTGGAG |
| 29 | 40 | GAGGTCGTCAAATTCTGACAACGATCAACTGGAGATAAAT |
| 30 | 40 | CCGATTGTTGTTATGTAACTCGCCTTACCCAGACCCTTAT |
| 31 | 40 | AAAGGAATCGGCGTGACACCACGATGATAAAGTGGTCTCG |
| 32 | 38 | GATGAACGTGACAGATCGCTGAGATCCTAAAGTTCTGC |
| 33 | 37 | AACAACGTAATAGACACTGGGGCCAGATGGTCCTCCC |
| 34 | 37 | GCTAACCGTGCACGGAAAAAGGAAGAGTATTTTCAAC |
| 35 | 25 | AGCGGTCGAGCGTGTGCAGGACCAC |
| 36 | 25 | ATAATGGTAACCCTTCTCAACAGCG |
| 37 | 21 | CCGGTACGCTGTTACCACCAC |
| 38 | 21 | GACAATTTCTTTTCAGGTGGC |

**Table S4**. Staple sequence for the pentagonal DNA origami of 84-bp edge-length. The sequence is represented by colors; unpaired nucleotides with blue, crossovers with orange, and the 14-nt seed dsDNA domain with green.

| Staple ID | Length (bp) | Staple sequences |
|---|---|---|
| 1 | 58 | CATCTTTCGATAGCACGAGGGTAGCAACGGTGAGGCTTTGAGTAGCTCTTTTTTAAAT |

| | | |
|---|---|---|
| 2 | 58 | ATTAAAGTCCAGACCTAAACAACTTTCAACTCAGCGGAGTGAACGCAGTTAGCAAACG |
| 3 | 55 | CTTTTGAAGGACAGAACCGCCACCCTCACTTGAGCTAATGCCACTACGATGTATC |
| 4 | 50 | ATCGCCAGAGGTCATAAATACGCATTTGGGAATTAGCCCTCAGATGAACG |
| 5 | 50 | TCATATTCTGTATGGATACATAAGGCTTTTGCAAAACTAATGCAATCCTC |
| 6 | 48 | GCGCCAAAGAGATTGGCATTACGAGGCATAGAGACGACAACGCAAAGA |
| 7 | 46 | GCCGCCAGACGTTCATAAGGGAACCCGCGACCTGTTTCTCCATGTT |
| 8 | 46 | TTGCCCTAAAAGATTGCGAACGAGTAGATGAGGCTGAGAGGGTTGA |
| 9 | 46 | GTGTCGAAATCGAACTGCACCCTCAGAGCCGGTGAATTAACCTAAA |
| 10 | 45 | GTTTATCTACTTTTAGTAGCATTCAGAAGGAAAAAAGGACCGTAA |
| 11 | 45 | TAGAAAATACGATTTTGGTTAGTAAATGAATACAAATAAGATACA |
| 12 | 45 | GTCGCTTTAGTTTATTAGATACATTTCGCAAAACAACCCTCAGGA |
| 13 | 45 | ATGCAACTAACATCGGAAGCACCGTAATCAGTTATTAGTTCATTA |
| 14 | 43 | CGTATTTATACTTAGGACGTTGGGTCCCCCTCAATTTATGCTT |
| 15 | 43 | TCTGAAGATTTTTACCACATTCAAGAAGTTTTGCTTTCAGAGG |
| 16 | 43 | ACCAGATCAAGTTCTTGACAAGAAACCAGACCGGTTTAAGCAA |
| 17 | 43 | GGGGTAACACCTTAGTAGTAAATTGAAGCAAAGCTTTGGATTG |
| 18 | 42 | CACTGAGTAATAAGATCTACGTTAATAATCGTCATGAAACGC |
| 19 | 42 | TCAACCGGGAGGTTAACACTATCCGGAACGAGGCGCGCCAGC |
| 20 | 42 | CTCCAAAAACCGAGAAATATTCATTGAAAAGAAAATTTTAAC |
| 21 | 42 | GGTTTAGTTAAGAGAGATGGTTTAATTTTTACCCTATAACCT |
| 22 | 42 | CCACCCTCCCTGCCGCGATTTTAAGAACAGTTCAGAAGGTGG |
| 23 | 42 | ATTGACAATTGAGGTCATCTTTGATAAGTTTATTTGCGACAT |
| 24 | 42 | AGTTAAACCAATTCTAAGAGGAAGCCCGTCAGTGAGCGCGTT |
| 25 | 42 | AGTTTGCGCATTTTACAAAGCTGCTCATAAAGACTATAACAG |
| 26 | 42 | TCGCGTTCAACGTACGGTCATAGCCCCCTAGCGACGCAGCGA |
| 27 | 42 | CCCAAATTTAATTCTGTCTGGAAGTTTCACCCTCAAGAATCA |
| 28 | 42 | AATAATATTCACGTACCAGTACAAACTAGATGATATAACGGA |
| 29 | 42 | AAGACAGAGTACGGGAGCTTCAAAGCGACCGGATACGTTTGC |
| 30 | 42 | GTTTAGCGATAGTTACCCTCAGAACCGCTTATTCTAATCATT |
| 31 | 42 | TTGATTCGGCCGCTGTCAGACTGATAAGTGCCGTCTGCAGGG |
| 32 | 42 | GTGAATTCATAAATTCATTTGGGGCGCGTTTCTTAAGAACCG |
| 33 | 42 | CCAGTAGCCGCCTCCGCATAGGCTGGCTAACAGGTTGAATAT |
| 34 | 42 | TCATGAGTAATTGCCAGGATTAGAGAGTGACCAGGCCTCAGA |
| 35 | 42 | GTGTACAACCTTTAGATGGCTTAGAGCTGAAGTTTAAAATCA |
| 36 | 42 | GCCGCCAAGCCAGCCCATTAAACGGGTATTTTGCGATTGCTC |
| 37 | 42 | CAATGACATGGTCAGACTATTATAGTCAGGGCTTGGCTGAGA |
| 38 | 42 | AGGTGAAAGCTGAAAAAACGAGAATGACACCTTATTATTTCG |
| 39 | 42 | GTCAGACCAAAAGGTGTCACAATCAATATAAAAGAGATAAAA |
| 40 | 42 | GAACCTACACCCTCAACAGCTTGATACCTATATTTCAAAAAT |
| 41 | 42 | TTCATCGCTTTAGCTTTGCGGGATCGTCATTCCATTCAAATA |
| 42 | 42 | CAGGTCTCAACTTTGAAACATGAAAGTATACCGCCGCGCCGA |
| 43 | 42 | TTCCACATAAGCGTGGTAGAAAGATTCAATAGTAATAAGACT |
| 44 | 42 | AACTAAAGCATGATAATGTTTAGACTGGTATTACACATACAT |
| 45 | 42 | ACAACATATAGCGTACCCAAAAGAACTGGGAATTGTGTAGCA |
| 46 | 42 | GGCTTTTCAACGCCCGAATAATAATTTTACGGAATCCAATAC |
| 47 | 42 | TGCGGAAAACGAACCAGGAGTGTACTGGTTTCGTCTGAAAAT |
| 48 | 41 | AATATTGACGTTTTATTCATTAAAGCCACCAGAATACCAGA |
| 49 | 41 | CGTAACGATCTTTTTTTGTCGTCTTCCAGAATGGTGCAGTC |
| 50 | 41 | TTTCAGGGATTTTGCCCAATAGGAAGTGCCTTGATAGTGCC |
| 51 | 41 | TATAAGTATATTTGAATAGGTGTATAGAGAAGGATATTAGC |
| 52 | 41 | GGCCGGAAACTTTCAATGAAACCATTCATAATCATACCGGA |
| 53 | 41 | ACTTAGCATCCCCAGCGATTATAAAAACACGAGGGAAGGTA |
| 54 | 40 | ACTCCGACCTTCAGCCACCACCGGAACACCATTGACTTTT |
| 55 | 40 | CATCAGACGAGATTTGCTCAGTACCAGGCGGTAATAAGGC |
| 56 | 40 | GGGTATCAGTTGATTTACCGTTCCAGGACAGCCAAGGAAC |
| 57 | 40 | TAAACTGGCTCAAAACAGTTAATGCCCAGAGCCGCTTTCG |
| 58 | 39 | TCACCGAGAGCCACACCAACTTTGAAAGTATGATAAATT |
| 59 | 39 | CACCACGGAACAACCCTCGTTTACCTAAGAGCGAGGCAG |
| 60 | 39 | ACCAAAAAAAAGGACTTGATATTCACAATTGGTTTACCA |

| | | |
|---|---|---|
| 61 | 38 | ACGAAAGATAAGAATACACTCCAAGCTTCAAAGTACAA |
| 62 | 37 | GGGGTCACCCATGTCTCCAAAAGGAGCCTTTGTATCG |
| 63 | 37 | CTCCTCACACCGTAATCGCCCACGCATAACTTATTCG |
| 64 | 34 | TAACGCCTAGCGAGAAGGTGGCAACATAGAAAAT |
| 65 | 25 | AATGCTGGACTAAAACCATTAGCAA |
| 66 | 25 | CCTTATTGAATAGACTCATAGTTAG |
| 67 | 25 | CATCAATTCAGCTTACCACCCTCAT |
| 68 | 22 | CGGAGATTAGGCACCATCACCG |

**Table S5**. Staple sequence for the pentagonal DNA origami of 126-bp edge-length. The sequence is represented by colors; unpaired nucleotides with blue, crossovers with orange, and the 14-nt seed dsDNA domain with green.

| Staple ID | Length (bp) | Staple sequences |
|---|---|---|
| 1 | 60 | TTTCGGAAATATCGTATGTTAGCAAACGAACGCAAGTTAGTAAATGAATGTGAAACATGA |
| 2 | 58 | AGAAACGTAAGCAGTTCAACAGTTTCAGCGACGATATAGCGTCCAATACTGTTCGTCA |
| 3 | 58 | TAGATAACAAGTACACACTGAGTTTCGTAGTATAGTGATACAGGAGTGTACTATAAGT |
| 4 | 58 | AAACAGCAAACGAAAGTAGCACCATTACCATAAGGCCGGAAAAAAATCTTTCTGAGAG |
| 5 | 55 | AATCAAATAGTTAGCGTAACAGGATTACGTATAAACAGTTACTTCAAATTAAGAC |
| 6 | 55 | CGAAATAACAAATAAATCCTCATGTTTGTAATTCTGTCCAGCGCCCAAACCCTCA |
| 7 | 55 | GGGCGACGATAACCAGCCTTTAATTGTAATCAGTTACCGACTTGAGCCATTATAT |
| 8 | 55 | AAAAGAATTTCTGTAAGGAAACCGAGGATAGAAAACCCGAAAGACTTCAACCTAT |
| 9 | 50 | AACTATTCAACCGATCACCGTCGAGATTTAGGAATACAAAAGGCACAAGA |
| 10 | 50 | ATTACCGACGACGAATGCTGTAGCTCAACATTAAATCAGAACCGTTGTGT |
| 11 | 50 | TATTCTTTTGCCAGAGAGGAAGTACATACATAAAGGTTACCAGATGGGAT |
| 12 | 50 | ATTCGAGATGCCCCGACTCCTCAAGAGAGATCTAAAACTGGCATGAAACC |
| 13 | 46 | ATCCGGTAGAATATGTGTCTGGAAGTTTGGTCAGATTAGCCGGAAC |
| 14 | 46 | GCAATAGTTATTTTAAAAATCAGGTCTTATTCATTTAGTAAGAGCA |
| 15 | 46 | CTGTCTTATCCCATCTCCAACAGGTCAGCGGGGTCGTACCAGGCGG |
| 16 | 45 | ATCAATTCTACTTTAGTAGCATTGAATAACCTAGCCCCTGACGAG |
| 17 | 45 | CCTTGATAAGTACGAAAGTACCGACAAAGATATAGTGTATCATCG |
| 18 | 45 | AAGTATTAAGTCCAGACTAATAACGGAATACTACGCAGCGTTTTA |
| 19 | 45 | TGCTCCTTTTCTTTTGACCCGGAATAGGTGTTACCGTACGCACTC |
| 20 | 45 | GAAGCAAAGCAGACTGGAAAAACCAAAATAGAACAACTATAGCCG |
| 21 | 45 | ACTACCTTTTAATCACCCTAACGGAACAACAGAGGTGAAAAGTCA |
| 22 | 45 | AGGCACATTCCTTCAGTTGATTCCCAATTCTCACCAGATGACCAA |
| 23 | 44 | TGATGCAAAATTGAGCATCAGCTTGCTTTCTTATTACATTAGAG |
| 24 | 43 | TTTGACTAATTTTTTACAAAATAAAGAAAAAGCCTTTTGTTTA |
| 25 | 43 | GACAAGACGGGTTTAACTGAACACGCGAGAAAACTTTTTTTTC |
| 26 | 43 | GCGCGGAACGCTTTTTTTAGCGAACCATTTTCGAGTTTCCAGTA |
| 27 | 43 | CCTGTCATTCCTTGGGTATTAAACGTCCTGAACATTTAGAAAA |
| 28 | 43 | ACAACACCGAATTTTTTAAGAAAAGCAAAGACACCTTTACGGAA |
| 29 | 42 | CGATTTTATAAGGCAATTTTCCCTTAGATAGCGTCGGTTTAA |
| 30 | 42 | TTTTGCGTTGAGATAGACTGTAGCGCGTATTAATTGTTAAAT |
| 31 | 42 | AAGAATAATCCAAACACCCTCAGCAGCGAGAACGAGGCATTT |
| 32 | 42 | AAACACCAAAGACATATTATTTATCCCAAACACCGTGTAAAT |
| 33 | 42 | TCGGTCATTGCTTCGAATCATAATTACTACAGCCAGCATCGG |
| 34 | 42 | ACGTTGAACATAACAACAGTTCAGAAATTATTCATCAACTAA |
| 35 | 42 | GAACCGCAATGGTCGGGCTTAATTGAGAGTTGCTAAAGGCAC |
| 36 | 42 | AAGATTAATCGCCAAGATACATTTCGCACACCCTCGGCGCAT |
| 37 | 42 | GCAACTAATTCACACCGCGACCTGCTCCGGAGATTAAGGCTT |
| 38 | 42 | AATTGCTCGCAGTCCCACCCTCAGAACCCACCCTCAGGAATC |
| 39 | 42 | GAACCACGCGAACGAATTTAGGCAGAGGCTCCCGAAACACTC |

| | | |
|---|---|---|
| 40 | 42 | AGACCGGCAGTGCCGGATTAGCGGGGTTAGCCCTCTAATCGG |
| 41 | 42 | AATGACCTGCTTTAGCCAAAAGGAATTAATAATAATGAAATA |
| 42 | 42 | CAACCTACAGACCAAGAGCCACCACCCTGACCATTTATTTAA |
| 43 | 42 | AGCGATACGACAGAATTGTGAATTACCTTCGGTCGGCCTTTA |
| 44 | 42 | CGTCGCTTTTCATCGTAGTAAATTGGGCGGATCGTTAAGAAA |
| 45 | 42 | CAACGCCTGAAGCCAGAGGCAAAAGAATAGAGGACGCCACCA |
| 46 | 42 | TGTTTAGAGCCGCCTCATCAAGAGTAATAACGGGTTATCCTG |
| 47 | 42 | TTAGTTTCAGAGCCAGATGAACGGTGTAAAACGAATTAAATC |
| 48 | 42 | AGGCTGGCACTACGTTTTGCACCCAGCTAACAGTAAATAACC |
| 49 | 42 | GAGCCACGCCACCCGCCAGAATGGAAAGGAATATACAATAAA |
| 50 | 42 | CAACATGTCATCGTATTTTCAGGGATAGAGTACCGTCTGAAT |
| 51 | 42 | GAGGTTTCAAGCCCAGCCGTTTTTATTTTTCAGCTAGAGCTT |
| 52 | 42 | TTACCGTATGGCTTAATGCAGAACGCGCACAAGCAAATAGGA |
| 53 | 42 | ATCGAGACTGTTTAGGTCATTTTTGCGGTCCAGTATACTCAG |
| 54 | 42 | ACCCATGATCACCGAGCGTCATACATGGGATAAGATCAACAA |
| 55 | 42 | AACGCTCACAATTTAAAATACGTAATGCCTGACCTACCCTCA |
| 56 | 42 | TGAGTAAAGCAAACCTAATTTACGAGCATGTAGAGCGAACC |
| 57 | 42 | TCCTTATCCAAAAGAGTTTTGTCGTCTTAGGCTGACTGCCTA |
| 58 | 42 | AACAAAGTGGCAACATCAAAAAGATTAAGGGGGTACTTTTGC |
| 59 | 42 | TTTGCTACGAGAGGATAGTAAAATGTTTGGATTGCATATAAA |
| 60 | 42 | GATATTCTTCATGATAACGAGCGTCTTTATATGCGAAAGGTG |
| 61 | 42 | CCTCAAAATAAATCGTCACAATCAATAGGAAACAATTTTTTC |
| 62 | 42 | TGCAGATAAATCTCAATAATAAGAGCAAAAAATTCAAACGAG |
| 63 | 42 | GAGCCACGAGCTGATTATACAAATTCTTCCAACGCGGAAGTT |
| 64 | 42 | CTTTGAAACACTAACTTGCGGGAGGTTTAACATGTAGTAGAT |
| 65 | 42 | AAGGCTCCCACATTTAAAGGTGAATTATTGAGGGAGACAAAA |
| 66 | 42 | AATCTTAACCAGTATTCATTTGGGGCGCCACCGGAAGAACCG |
| 67 | 42 | GAGGGTATCCAATCCGGCTTAGGTTGGGTTGGGAAGGTAGA |
| 68 | 42 | CCAGCAATAACCTCGCAAGACAAAGAACCCTGAACATTTCTT |
| 69 | 42 | TCCATTACTTGACAACCGCCTCCCTCAGCTATATTTAAAGCC |
| 70 | 42 | AACTGGCGCCCACGTAAAAACAGGGAAGTATTTTAATAGTGA |
| 71 | 42 | CCATCGAAGAGTCAGTTAATTTCATCTTAATAACACATAACC |
| 72 | 42 | CAGAGAGCTGACCTTTAAGACGCTGAGATAGCAGCTTTTAAG |
| 73 | 42 | GATATATTATGCGAACCGTAATCAGTAGGCTTAGAAAATTTA |
| 74 | 42 | ATGGTTTAATAGCACTGAGGCTTGCAGGTTTAATCATCAAGT |
| 75 | 42 | TTTCAACGAGTTAAACGTCAAAAATGAAGAAATACAAAACAT |
| 76 | 42 | TTGCCTTATCCTTGCGACCGTGTGATAATTGTTTAAGGCCGC |
| 77 | 41 | ATAAGTGCCGTTTAGGGTTGATATACACCAGTACTACAACG |
| 78 | 41 | ACACTATCATTTTTCGTTTACCAGACGGAGTGAGTAAAGGA |
| 79 | 41 | CGTTGGGAAGTTTTCTACGTTAATATTGATACCGTTGCGCC |
| 80 | 41 | ACAAAGCTGCTTTCAGTGAATAAGGGTAGCAACGTAGAGGC |
| 81 | 41 | GAGGCGCAGATTTAATCATAAGGGAACCCCCAGCTTACCAA |
| 82 | 40 | ATAAGATTCTAAAAACAAAGTACAACATGTTACCGATTGG |
| 83 | 40 | ATAATTCCTTATAGCATTCCACAGACTTGCTCAAGTGCCT |
| 84 | 40 | TAAGTCTATCTTTAAAGGAATTGCGACGAGGCAGAATCCC |
| 85 | 40 | AATATTGACGGAATATGGTTTACCAGTAAGCCCCAAAAAA |
| 86 | 40 | AAGATTCTCGGTTTGCTAATATCAGAGAATATGTAAATGC |
| 87 | 40 | AAATACGCATTATGACAACAACCATCTCATTATAATGAAA |
| 88 | 40 | GTATCCCAGAGCGGACTAAAGACTTTATTACCCGGAACCA |
| 89 | 39 | CCTGATAAACCTAGCAAGCAAATCAAGGTAAATAAATAT |
| 90 | 37 | AACGAGGCTTGCCCCTTATTAGCGTTTGCCTTTCATA |
| 91 | 37 | ATCTTTGACCGAACGCCGCCGCCAGCATTGTAGGTTG |
| 92 | 25 | ATTTATCCGTCACCACCAGTCAGGA |
| 93 | 25 | GCATCAAAATCACCAAATCAACGTA |
| 94 | 23 | ATTGAGTCGCCAAAGGGAAGGTA |
| 95 | 21 | TTTAAGATTAGTTCCTTTAAT |
| 96 | 21 | TAAATTACCCTTTTATAGTCA |

**Table S6**. Staple sequence for the pentagonal DNA origami of 168-bp edge-length. The sequence is represented by colors; unpaired nucleotides with blue, crossovers with orange, and the 14-nt seed dsDNA domain with green.

| Staple ID | Length (bp) | Staple sequences |
|---|---|---|
| 1 | 60 | TCTAAAAACCATTATAAGAACTGGCTCAAATTGGGCCCATCCTAATTTAATCACTAACAA |
| 2 | 58 | AGAGACTATTTAACTTGCCATCTTTTCATATAATCACCGGAAGACAGGTTAGGCAGGT |
| 3 | 58 | GGTATTAAACAAAGTGGCAACATATAAAAGTCAAAGACACCAATGTTTTTTGCAACTA |
| 4 | 58 | AGAATAAGCGTAGAACCTGAAAGCGTAAGATTGGCACAGACATCACTTTTGTAGAAGA |
| 5 | 55 | CTGAAAGAATACCAAGTTACCGAGAAATCGTCACCCTCAGCCCCTGCCGAGCCAC |
| 6 | 55 | GTAACGGTTTATCAACAATAATTTGAGAACAGTTGAAAGGAATGGTCAGTGAATT |
| 7 | 55 | CATATTCGATAGCAGCACCGAATCAGGAAATATTCATTGAAACACTCAATTTTCC |
| 8 | 55 | TTTAAACGAGGCAAAAACATAGCGATAGGTGAATACTTTAGCGTCAGACATGGAA |
| 9 | 50 | TTCGCAAATTGAGGTCAATAGATAATACGATAAGTCTTTAATCAGTTAGC |
| 10 | 50 | CACTAAATCCCCCTGACCATAAATCAAATAATCAGGCTATTAATAACAGC |
| 11 | 50 | TAGGAGGATTAAGACCGAACGAAGTCAGGACGTTGGGAAACACGTAGAAA |
| 12 | 50 | AGCGCATTCAGAAAAAGTTTGCACCTTGCTTCTGTATCCTTGAAAGAATA |
| 13 | 50 | TAATGCCAGCGAAACAAGACAAAGAACGAAAATCGCGCCACCCTAATCGT |
| 14 | 46 | ATTTGTAATAGTAATCAAAAAGATTAAGGTAGCACCAAAAATGAAA |
| 15 | 46 | TTAGTAACCAGTACAAGGTGGCATCAATGAACCTCATTAAATCCTT |
| 16 | 45 | CGCCTGAATTTTAGTTAAAGGCCGCTTTTTATTCTTACCTACATT |
| 17 | 45 | CACCAAGGAAGTTAGACTTCAAATATCGCGTACTTGAGAGGGAAG |
| 18 | 45 | AAGTACGGTGATAAAGGTTACCAGAAGGAAATCATTCCGCTGCTC |
| 19 | 45 | CAGACGATTGTTAGCGTAATTTCATTTGAATAAATCATCCATTAA |
| 20 | 45 | CTAATAGATTATAATATCTTGAGATGGTTTAGCGATTTGATACAT |
| 21 | 45 | ACTCAAACTACCTTCTGTTTTCAGGTTTAACTAAGGCGACAATGA |
| 22 | 45 | TTGCTTCTACTTTTAGTAGCATTAGCTAAACAGCCAGCACATTAT |
| 23 | 45 | TGCAGAAAGACTTTTATCTGGTCAGTTGTATATTTTCCACAGACA |
| 24 | 43 | AGAAATTAAACTTATACCGATAGTGGATAAGTGCTTTCGTCGA |
| 25 | 43 | GAGAAGATATTTTCCAAATCAACGTTATTACAGGTTTTAGAAA |
| 26 | 43 | AGGTTAGGACTTTTTTTCATGAGTGGTAATAAGTTTTTTTAA |
| 27 | 43 | CAGACTTCTGTTTATTTTGCTAAACCGTAACACTTTTGAGTTT |
| 28 | 43 | TTTATCTGATATTGTCGAAATCCGGAAGTTTTGCTTTCAGAGG |
| 29 | 42 | AGCAATATTATTTTGAGTAATCTTGACAATCAGTTTGCTGTA |
| 30 | 42 | TTTGTCAAATATAAGAGATTTAGGAATATCATCAACATCGTA |
| 31 | 42 | CTGACCTCCACATTGAGCTTAATTGCTGCAATCAAATGAAAT |
| 32 | 42 | GGAATCAAGAAACATAGAAAATTCATATTGGCTTACAACTAA |
| 33 | 42 | TGCAGATGGCGCATGCCCAATAGCAAGCTTAAGCCCAGCGCC |
| 34 | 42 | CAGACGGAGACGACCCGGAAGCAAACTCTGACGGACAAAGTC |
| 35 | 42 | CCTCCCGCCCTGAAAATTATTCATTAAAAACCAGAGATAAAA |
| 36 | 42 | CAAAGCGAGGCCGGAAACGATTTTTTGTACGCTAAAACGGAG |
| 37 | 42 | AGCCAGATGTAGCGATCAATATATGTGACTTAGATAACCTAA |
| 38 | 42 | CGCATTACTTAAATCTCCATGTTACTTATAGCGAGCGAGCTT |
| 39 | 42 | CACCCTCTCAGAACCGCAGAGGCGAATTCCAATCGGACAGCA |
| 40 | 42 | GCCAGCCGACCAGTTTTACATCGGGAGATGACCTAGTCGCTG |
| 41 | 42 | GTCACCGTTTAATTAGGCTTTTGCAAAACGACCTGCAAGATT |
| 42 | 42 | ATCACGCGCCCTAAGGATTATACTTCTGTTACCAGTTAATTG |
| 43 | 42 | GTTTTTACAGAAGATTATCAGATGATGGGAATCGCTCACGTT |
| 44 | 42 | AGGGATTCTGCAACAAGAAACCACCAGAGGCATTTAATAGAA |
| 45 | 42 | GGGCGCGCAATCAAACAAACAATTCGACACATGTTCCAGACG |
| 46 | 42 | AATTCTGTCCTTATCCAAAAGAACTGGCCGAGCATCAGAACG |
| 47 | 42 | TTGCGGAGGTTTACCAATAATAAGAGCATTACCGCAGGCTGG |
| 48 | 42 | TAGAGAGTGAGGGACGCTAATATCAGAGTCCGGTACTTTGAA |
| 49 | 42 | CAAAGCGGGTGAATGAATTAACTGAACAACTTGCGCGAGGCG |

| | | |
|---|---|---|
| 50 | 42 | AATTGAGAAATCAGCGGTGTACAGACCAACATAACGTCATTT |
| 51 | 42 | ATTAGCAGATTGCAAATGTTTAGACTGGAAAGTACCGAGCGT |
| 52 | 42 | CCAATCCGAGCCTAACCAAGCGCGAAACATAGCGTTCAGAAG |
| 53 | 42 | CTTTCCAAAATAAGAAACGTCACCAATGATTATAGCCAATAC |
| 54 | 42 | TGCGGAACGATTATATTTGCCAGTTACAAAATATATCTTTGA |
| 55 | 42 | CCCCCAGTCGTCATTCTTTACCCTGACTAAACCATATTTATC |
| 56 | 42 | CTTAGAAAATCGTCTAGCGACAGAATCACGAGAATCAAATGC |
| 57 | 42 | AACGAAAAGGTCTCTGAATTTACCGTTCAGGCACCTAAGACG |
| 58 | 42 | GAAACAGGAGTCAATAATGCCACTACGACAGTAAGTCATTAA |
| 59 | 42 | CTGAGAATACATAACGTTTTCATCGGCATAAATCCCGTCATA |
| 60 | 42 | CATGGCTAAATACGTAGTGAATTTATCATACCTTTTCATAGC |
| 61 | 42 | ACGGGTATTTGATGTATTCACAAACAAATTTTCGGTTTAATG |
| 62 | 42 | CCCCTTAGCCTTGAATACAGGAGTGTACGAAGTTTAGGTCTG |
| 63 | 42 | AAAGACAATAAGAGGCCAAAAGGAATTAAGATGAAATATAGA |
| 64 | 42 | CCTGAGCTATGTAAGCAACGGCTACAGATCAGTGCCCGCCGC |
| 65 | 42 | GGAACCGACCAGAGCTTGAGTAACAGTGGAGGGTAATGCTGA |
| 66 | 42 | TCGGAACCCCGTATCCACCAGAACCACCCCTCCCTTCAATTA |
| 67 | 42 | TGCAAATATTCATTCAGAGCCGCCACCCAGAGCCGAAACAGT |
| 68 | 42 | AGAGGGTGGCGTTTTAAGGGAACCGAACCTATCATTCAGGAT |
| 69 | 42 | CAAATATTTGCTTTTGGATTATTTACATATGGAAAGAAACAT |
| 70 | 42 | GAAAGTACAGGGAGTTAATTTCATCTTCAACAATATTCACCA |
| 71 | 42 | AGGCTTGTTAAGAGCAGGAAAAACGCTCTGGCAGAACGGATT |
| 72 | 42 | GTCACACATTGCAAGCTGAGACTCCTCAATATTCGAATTTAA |
| 73 | 42 | CAGTAACAAATACCCACGCATAACCGATAGAGAAGTATTACC |
| 74 | 42 | TGGTTTGAGTACCTAATAAAAGGGACATAGAACAAGATTAGG |
| 75 | 42 | ATTAGCGCATCGCCGACCGTGTGATAAAGTCAGATAACAGAG |
| 76 | 42 | CAACAACGGGTTTTTGCTGGTAATATCCTCTGGCCGAATATA |
| 77 | 42 | ATAGAACTCGGCCTGCTCAGTACCAGGCTGCGCCGTTAAATA |
| 78 | 42 | AGAGGACCGAGGCAAATTGCTCCTTTTGAAAGGGCCCACAAG |
| 79 | 42 | TAGAACCTCATATGCTTGCTTTCGAGGTTTGATATGATTAGT |
| 80 | 42 | CTATTAGCTTCTTTAAGTATAGCCCGGATTATCAGCGTTATA |
| 81 | 42 | TATCGGTATAGGTGCCGTTGTAGCAATATCTTTAAGAAGGGT |
| 82 | 42 | CAAATTCAATAATGTGCGCGAACTGATAAAATTAATATCACC |
| 83 | 42 | GTACTCAGGAGCCTTATAAAGCCAACGCATTGTTTAACATCG |
| 84 | 42 | AATCCTGTCAACAGAAAAGGCTCCAAAAGGAGGTTTCTGTCC |
| 85 | 42 | CCATTAAAAAAGAGTAGTACCGCCACCCTCCAAAATAGGGCT |
| 86 | 42 | GAAAATCTCAGAACTGAGGCCACCGAGTAAATACCTCAATAT |
| 87 | 42 | TAATTGACAATTCAGAACGAACCACCAGTAATCAGCGCCACC |
| 88 | 42 | CTCAGAAAATTTTTCATATTTAACAACGTTCCTGATAAAACA |
| 89 | 42 | CATCATACCAACATGAATTGCGAATAATCCGCCACGAGAAGT |
| 90 | 42 | GAGGTGAGAATCCTCCTCAGAGCCACCAACTAAAGGTAATTT |
| 91 | 42 | AGGAACACCCTCATGGAACGGTACGCCAGGCGGTCGAATTAT |
| 92 | 42 | AGGCAGAAGGAGCGAGTATTAACACCGCTTAGACATTTCAGG |
| 93 | 42 | GATAGCAGAGTGAGTCGAGCCAGTAATACGGAACAAGTGCCA |
| 94 | 42 | CATTTTGAGAGAATCAACAGTTTCAGCGAGCCCAACGATTAA |
| 95 | 42 | CGCTGAGAGGAGGCTAGGAACCCATGTACAACTTTATAAAGT |
| 96 | 42 | AGGCTTAAGATAACGACATTCAACCGATTACCTTTTAGTAAG |
| 97 | 42 | AAACCCTAGCTGAAAAACTACAACGCCTCGTCTTTCAGCTAA |
| 98 | 42 | AAGTATTCGCGCCTATCTAAAGTTTTGTGTAGCATTCATTTG |
| 99 | 42 | TAAATATCAACAGGAACCCTCGTTTACCTCAATCATAGCGAA |
| 100 | 42 | ACCTTATATTTCAACCTGAACAAGAAAAAGAGCCGAAGGTTA |
| 101 | 42 | ACCAAAAGCCGGAAGGAGGTTTTGAAGCGACGGGATATCACC |
| 102 | 42 | TAATAACATAATCGGCTTGCCCTGACGAGAAGAAAGATTCCC |
| 103 | 42 | CCAATCAGGAATACTACGCAGTATGTTAAACAGTTAATCTAC |
| 104 | 42 | GTTAATAGAATAAGGCTGTCTTTCCTTACCGAGGATAGAAAA |
| 105 | 42 | ATTCAGTAAACGAAGTTTCATTCCATATGCAAACGAACGCAA |
| 106 | 42 | TACATACTCTGGAACTAACGGAACAACATAACAAAAAGAACG |
| 107 | 42 | AGCAACATGACCAATTCTAAGAACGCGAAATTGAGGGGAAGG |
| 108 | 41 | AAACATCAAGTTTAAAATTAATTACACCTTTTTATCGGCTT |

| | | |
|---|---|---|
| 109 | 41 | TGCACGTAAATTTAATAAAGAAATTACACCGGAATATTACT |
| 110 | 41 | TGCCCGAACGTTTAATTTTAAAAGTAAAGGTAAATTCTGTC |
| 111 | 41 | TTTTTAAGAATTTAGCAGATAGCCGAACCAAGTATCTCATC |
| 112 | 41 | ATAGCAGCCTTTTGAGAGAATAACAATTTTGCACTTACAAT |
| 113 | 40 | GGGTATCATCGCCCTGAATCTTACCATTAACGTCATTACC |
| 114 | 40 | CGGGGGGCTTTGGGGTTATATAACTAAAAAGAACACCACC |
| 115 | 40 | GAGGGGAATTTCAAGCCTGTTTAGTATACCATATGAATGG |
| 116 | 40 | CGTCAATGAATTGACGACAATAAACAAACTCGTAAATATC |
| 117 | 40 | GATTCAGAACCGCAAGCAAGCCGTTTGCTATCTAGTTTAT |
| 118 | 39 | TTGACGCTCCATATTTCGGAACCTATGCGGGAACTTTTT |
| 119 | 39 | GCCCTCATATTATAACCTGTTTAGCGCAAATCGATTTAG |
| 120 | 37 | ACCGACATTGAGTAAGCAAATGAAAAATCTTATCACC |
| 121 | 37 | AGTTGCTTAAAAACCCATTTGGGAATTAGATCAAAAT |
| 122 | 34 | AGTAGTATTATACCGTAGATTTAGTTTGTATCTT |
| 123 | 25 | CAGCATTCCAGAGCGATGATGAAAC |
| 124 | 25 | AATAACAATATTTTTCAAAATTATT |
| 125 | 25 | GCTCAACCGGAATATACCGAAGCCC |

**Table S7**. Staple sequence for the pentagonal DNA origami of 210-bp edge-length. The sequence is represented by colors; unpaired nucleotides with blue, crossovers with orange, and the 14-nt seed dsDNA domain with green.

| Staple ID | Length (bp) | Staple sequences |
|---|---|---|
| 1 | 58 | AGGTGCCGACCTGACGAGGAAACGCAATACTGGCTGTAAGAGCAACACTATTCCCTCG |
| 2 | 58 | TTATTAGACGGGTAGTCCAGACGACGACAATAACATGTTCAGCCAGCTTTGCACCGCT |
| 3 | 58 | TATTGACTTTGAAATAAATAAGGCGTTAAATATAAACACCGGTTTCCCTTTCCTTGAA |
| 4 | 55 | AGAAAACAAATCGCAGCAGCACCGTAATCCCCCAGAACGCCAACATGTAGGGCCT |
| 5 | 55 | ACAGGTAATAGCTATGATAGCCCTAAAATCAAAGGCTTCAAATATCGCGGGTAAT |
| 6 | 50 | CTTCGCAATTAATTAATTTAACCGATTATACCAAGCCATCGATGCAGAGG |
| 7 | 50 | TGAGCGAAGATTCATCGAAAGAGCGAAAAACCGTCTCGCGAACTCTTACC |
| 8 | 46 | CCAGCAGAAATCGGGGTCATTTTTGCGGTTAACGTCACTGAGTTTC |
| 9 | 46 | AAAGGGACTAAAGGAAACAGTTCAGAAACAGACGAACTTTAATCAT |
| 10 | 46 | AAATTAATTAGAATCGCTATTAATTAATAATCATACGCAGACGGTC |
| 11 | 45 | TCTGGTGCCGTAATTCTAAATACGTAATGCCCGGTCATAATATAC |
| 12 | 45 | AAGCGCACCAACAGGAGTGTTGTTCCAGAAACAGAACCCTCAGAA |
| 13 | 45 | AACATAGCGACGTGTGAGAGGACAGATGAACGGAGGGAACTTGCC |
| 14 | 45 | AGATATTCTACTTTAGTAGCATTGTCTGGCCAGGAATCCGAGGTG |
| 15 | 45 | GAGAGATAACGCAAGAAATTAAAAATACCGATAAAGAATCGAGCT |
| 16 | 45 | AAAATCAGGTAGGCATACATTATACCAGTCAAGGAAACAAGCGTA |
| 17 | 45 | TTTGTATGGCTTTTTAATTGCTGAATATAATTATTATTCACAGAC |
| 18 | 44 | GCTGGCGAATGCTTTGGACAGAATCAAGTTCACTAAACAGAGGC |
| 19 | 43 | TCAAAGCGTAGTTCAGGTTTAACGTAAAACGACGTTTGCCAGT |
| 20 | 43 | CCGTTTGCCCGTTATTAATTTTAAGCTCACTGCCTTTCGCTTT |
| 21 | 43 | AACTGGCCAACTTAGAACCCTTCTGTAAAGCACTTTTAAATCG |
| 22 | 43 | GAATTAGCAATTTTTGATTAGTAAACGTATAACGTTTTGCTTT |
| 23 | 43 | TCGAGAAAATCTTATCACCTTGCTTCCTGTTTGATTTTGGTGG |
| 24 | 42 | GGCCAACTAGAAGTCTGGTAATAAGTTTAGGAATTCCCGACT |
| 25 | 42 | CAACTAATAACGGGTACATTTGAGGATTGCGCGGGAATAACC |
| 26 | 42 | AACGGAGCGGAAACCAATTACCTATCCTGAGAAGTACCATTA |
| 27 | 42 | TGCGGGAAATGGTCGAGAGGCGGTTTGCTAGATAAGTCAGTG |
| 28 | 42 | CCGTCAAGTATTGGAGATACATTTCGCAGGTTTTGAAAGGAA |
| 29 | 42 | CCTTGAGAGAATAGAAGCCTTAAATCAAGACCATTGCGCCAG |
| 30 | 42 | TATAAAATGAATAAAAATGTTTAAAATGCTGATGCAGCTGCT |

| | | |
|---|---|---|
| 31 | 42 | TCAGTTGTGCAGCAAACTAAAGTACGGTTCTTTCCTCGTCTT |
| 32 | 42 | GGCTGAGAGTTTTGAGAGCCTAATTTGCAATATGCAGCGGTC |
| 33 | 42 | TCCTTTTGCAGCCTATAGGAACCCATGTTATAGCCCTGAGAG |
| 34 | 42 | GCAAACTTTAGACGCCCTCAGAGCCACCTACCGCCGGTGAGG |
| 35 | 42 | TCAAAGCACACCCTAATTGAGTTTCAGAACCGCCAGGAGAAT |
| 36 | 42 | ATTGCATACATTCATAAAACGAACTAACTTTTTAAATTAGTC |
| 37 | 42 | ACAGCCAGCTGTAGCCAGCAGGCGAAAAGAACCTCGGGGTTT |
| 38 | 42 | CCCTCAAATAGCGAGCTTGAGATGGTTTCCTTATTAGTAATA |
| 39 | 42 | GCGTCCAAATAGTAGGCTTGCCCTGACGTAAAGGTAAATGGA |
| 40 | 42 | CTCCGGCGACAAAGTTACCCAAATCAACATAAGTTAACGCTC |
| 41 | 42 | GAATTTAATTTCATTGACCTTCATCAAGTACCAGCAATATCC |
| 42 | 42 | CTTCTGTCCTGTTTTGTTACTTAGCCGGCTTGAGCATCACGC |
| 43 | 42 | GGAAACAAAAGCCATCATCGCCTGATAACACCATTGTTTTTA |
| 44 | 42 | CGCAACTAGCCAGTGAGGCAAAAGAATATGCCTTTCGGATTC |
| 45 | 42 | CACGCTGCAATCAAAAGAGAAGGATTAGATAGTTAAAAATAA |
| 46 | 42 | AGTTTGAGAACAAGAGACAGCATCGGAACCTCAGACATATCA |
| 47 | 42 | TCACGTTGAAACCAGTTAAAGGCCGCTTCACCCTCGTTTGGA |
| 48 | 42 | TCTCCGTAACGGGTCCACGCATAACCGACAGCATTCTGATTA |
| 49 | 42 | TTTCATCAGCCGTTCTTGATACCGATAGGATATTCAACAAAG |
| 50 | 42 | AGCCCTCGATTAGCAAATATCAAACCCTGTTTGCCCTCAACA |
| 51 | 42 | TGTTTAGCGAACCTGCGAATAATAATTTGAGTGTAATTAGAC |
| 52 | 42 | TTAGTTTGATTAGTGTTTCAGCGGAGTGTAACAGTATTAGAG |
| 53 | 42 | TTCCATATCTTACCAATGAATTTTCTGTGGAACCTAAGGTTA |
| 54 | 42 | TGTTTTACAGTTACGCGTAACGATCTAAACTCCTCTATCTGG |
| 55 | 42 | GGTGGTTCTAATAGGCCCGTATAAACAGTTCAACATGCTATT |
| 56 | 42 | GAAAATAGATAAGACAAAATCCCTTATATGCCACGCGGAATA |
| 57 | 42 | GGATAGCCACCGTAACCGCCTGCAACAGAATCAAATAATTGC |
| 58 | 42 | GGTGTATAAGCCCATTACAGAGAGAATAGTACCTTAGAATAG |
| 59 | 42 | CCCGAGATATTAACCTCAGGAGGTTTAGACCCTCAAACAGGG |
| 60 | 42 | CGGTCAGTAGGGTTGTCAGGATTAGAGAACATAAATTTTCAG |
| 61 | 42 | TCCAGACTGAAAGTAACAGTTGAAAGGAACCGCCTAGTTTCA |
| 62 | 42 | TAACTGAGAACCAGCAAGAGTCCACTATACGAACCGCCCAAT |
| 63 | 42 | AATAAGACCACAAGGAACAAAGTCAGAGTTTTAATCGTGGAC |
| 64 | 42 | ACGAGCGGTCTGGAGGCCCTGAGAGAGTGCAAATCATTAAGA |
| 65 | 42 | TTTAATGATCAGGGATTAAGAGGAAGCCCAGTTGACATTATT |
| 66 | 42 | GAAGCCCGGAACAAGATTTAGGAATACCCAAAAAGCGATGGC |
| 67 | 42 | CCACTACAATGGCTGAAAAGTAAGCAGAACGTTAAACTAATG |
| 68 | 42 | AAAATCTTAGCCGAAGACAATATTTTTGGTGAACCAAAGCGG |
| 69 | 42 | CAGATACCAGAAGCATCACCCAAATCAAGTGGCACACAAAGT |
| 70 | 42 | AGAATACGTTTTTTCTGACTATTATAGTATAACGCGGGAAGA |
| 71 | 42 | TACCAGAGGACGTTCAAAAGGAATTACGCTTTACCGGGGTCG |
| 72 | 42 | AACAACTTTAATGCAGGAGCACTAACAATTTCTTTAGTAGAT |
| 73 | 42 | AACCAAAATGCTTTGAGCCCCCGATTTACACGACCACGCAGT |
| 74 | 42 | GAGTAGTCAAACGTAGATTCACCAGTCAGAGCTTGTGAATCC |
| 75 | 42 | ATGTTAGAAATTGGGAGGCTTTTGCAAATATTCATACGGGGA |
| 76 | 42 | AAGCCGGCATTGGCAGAAAATACATACAAGAAACATTGCCAG |
| 77 | 42 | TTATTTACGAACGTGGAATCGTCATAAAAGAAGTTCCAGAAC |
| 78 | 42 | AGGGGGTATACTGCGGCGAGAAAGGAAGTCGTCTGGGCAACA |
| 79 | 42 | CATTCAGGAAACGCATTTTGACGCTCAAGGAAGAACTGGATA |
| 80 | 42 | GGAGCGGTACCTACAAAGACACCACGGAGTAACAAAAATCCA |
| 81 | 42 | ATGGAAAGCGCTAGGGGTTATATAACTATATGTGAAGCGAAA |
| 82 | 42 | ATCGCAATTAGGTTGGCGCTGGCAAGTGCAGGAAATATTTTG |
| 83 | 42 | GAACCGGCAATAGAGCCAGCCATTGCAATAGCGGTTTTTAAC |
| 84 | 42 | TCACAATATATTCAAACGCGAGAAAACTACTACCTCACGCTG |
| 85 | 42 | CGCGTAATATTACCAAATTCATATGGTTAGTAATCATATATT |
| 86 | 42 | AGAACAACCACCACCATAGGTCTGAGAGTTTTCAATTGACAA |
| 87 | 42 | TTAGTTATCAAAATACCCGCCGCGCTTATGCTGGTGCCAAAG |
| 88 | 42 | GCGCATAGGCGACACAAACTATCGGCCTATGCGCCCAATAGT |
| 89 | 42 | ACAAAAGGGCTGGCCTTCTGACCTAAATGAAGAGTGCTACAG |

| | | |
|---|---|---|
| 90 | 42 | GGCGCGTAAGAACTTTCAACCGATTGAGGGTGTACTTTGAAA |
| 91 | 42 | TGAGTAGACTATGGGATTAAGACGCTGATTAATGGAGACCAG |
| 92 | 42 | TACCGACTAGCTTATTGCTTTGACGAGCTAACATCAGGTAAA |
| 93 | 42 | TTGCACCGCGAACGTCACCAGTGAGACGTATCTTTCCCCTGC |
| 94 | 42 | GAAAAAGAAATCGTCAGAGCGGGAGCTATCTGTCCCATTTGG |
| 95 | 42 | CGCGACCAGCCAGCACCGAGTAAAAGAGAACAGGAAACCTTG |
| 96 | 42 | GAATTAGTGCTCCAAGTATCATATGCGTAGTGAATGGCCGAT |
| 97 | 42 | TAAAGGGTGAGGCCAAAATCACCAGTAGATTGTGTATTCTTA |
| 98 | 42 | TAATCAGATTTTAGAATCAATATATGTGTATACAACGAAATC |
| 99 | 42 | CCAGTATGTACATAACAGGAACGGTACGCCAGAGATTTTAAT |
| 100 | 42 | GCAAGGCATTTGTAACGCTCAACAGTAGTTACCTTGCAAAAG |
| 101 | 42 | AAGATGATTCATTTGTCACCAATGAAACGCGAAACTTGAGAA |
| 102 | 42 | CGAATTATGAAACACAATTTCATTTGAAGGCTTAAAAAGTAC |
| 103 | 42 | TCTAAAAGGCAACATGATTCCCAATTCTCAGCTACTTTGCTA |
| 104 | 42 | GCCTGATAGGGGGAAGGGCGATCGGTGCATTTAGGACACTCA |
| 105 | 42 | ATTTTCGGTTGGGATGTGCTGCAAGGCGACAATAAAGCGTCA |
| 106 | 42 | AACCTAAGCGCGTTTTACATCGGGAGAAATTAAGTCAGGCTG |
| 107 | 42 | GACTGTAAACGAAAAATAAGAGAATATACGCCATTTGGGTAA |
| 108 | 42 | CGCCAGGGTACCTTTTCATCGGCATTTTACTACGAGACAAAA |
| 109 | 42 | AGTAACAGTTTTCCGGCAAAGCGCCATTAAGTACCAGGCACC |
| 110 | 42 | GGTAAAGGAAACCACAGTCACGACGTTGTCAGATGAGCCCCC |
| 111 | 42 | CTATTTCATGGGATAATTTTATCCTGAATAACAGTGCTGATT |
| 112 | 42 | GCTACAGAGCCACCTAAAACAGAAATAAGCTTGCAAGATCGC |
| 113 | 42 | GCCTGTTCTCAGGATGCCTGCAGGTCGATTGCACGACCGGAA |
| 114 | 42 | AAATTATCTCTAGAACGACAGTATCGGCTATCAACAGCAACG |
| 115 | 42 | CCGCCTCCGAGGGTAATAGATAAGTCCTGGGGACGGGATCCC |
| 116 | 42 | CGGGTACAACCTACGCCGCCACCCTCAGCAGCGAAAAAAATA |
| 117 | 42 | CCCTCAGAACCGCCAATGGAAGGGTTAGCGAGCTCATCTGCC |
| 118 | 42 | ATATCCCACCGTGCGAATTCGTAATCATTCTGAATACCCTCA |
| 119 | 42 | TTATACTGGTCATAATGGGCGCATCGTAATCCTAAATCGTCA |
| 120 | 42 | GAGCCACTTGCGGGTTTACGAGCATGTAGGTGTAGGCTGTTT |
| 121 | 42 | CCTGTGTCCTGATTAGAGCCGCCACCAGGCAGGGAATCAATA |
| 122 | 42 | GAGGCTTAACCACCTTCATCAATATAATGAAATTGGGATAGG |
| 123 | 42 | ATCGGCTCGTAATGTTATCCGCTCACAAATGGCAAACCAGAG |
| 124 | 42 | TCAGATGTTCCACAAACGGCGGATTGACGTCTTTCGGTCGCT |
| 125 | 42 | CCGCCGCTATATTCCTTATCATTCCAAGGGGAACACAACATA |
| 126 | 42 | CGAGCCGCATATTCGACAGGAGGTTGAGCCATCGCATTAAAC |
| 127 | 42 | ACAACAAGCAGGTCAGCGGAATTATCATGAAGCATGTCGGAT |
| 128 | 42 | CAAGTACACAACCCAAAGTGTAAAGCCTCAGAAGGAGACGAT |
| 129 | 42 | AAACCACGGGGTGCAATGTGAGCGAGTACGCACTCGACAATG |
| 130 | 42 | TGGCCTTTTGCGCCATCGAGAACAAGCAAACATTACTAATGA |
| 131 | 42 | GTGAGCTTTTGCGGACAAACAAATAAATTAAACAGTTTATTT |
| 132 | 42 | AATTTCTCCTCATTAGTAACATTATCATAACTCACAGCCAGC |
| 133 | 42 | TCATCGTTTCCTGTATTAATTGCGTTGCAAGTTTGAAAGCCA |
| 134 | 42 | GCCCTTCATTGAGGATTATTCTGAAACAGTTAGTAAACGCTA |
| 135 | 42 | TCTCCAAACATGGCGACAACTCGTATTACGGGAAAAAAGGTG |
| 136 | 42 | GGTATTCGAGCTGACCTGTCGTGCCAGCACAATTCTTTTGAT |
| 137 | 42 | TTTACAATGCATTATTCATTTGGGGCGCTAAGAACTTGAAAA |
| 138 | 42 | GATACAGTTTCACGGCGAGGCGTTTTAGCTATATTATGAATC |
| 139 | 41 | TGTGAATTACTTTGCGATTTTAAGAAATAACGGATCAAAAG |
| 140 | 41 | AATCATAAGGTTTGAACTGACCAACGGAAATTATTAAAGGT |
| 141 | 41 | ACTTTTTCATTTTAGTTTCCATTAACGTTTGCCATTCATAA |
| 142 | 41 | AGCCTTTAATTTTCGGTTTATCAGCAAGCGCAGTTAATTTA |
| 143 | 41 | GTCACCAGTATTTTACAACGCCTGTTACCAGGCGTGTGCCG |
| 144 | 40 | TTCCGCAAATGAAGGGTTGATATAAGACCGTAACAAAAAT |
| 145 | 40 | GAACCCATTCTGGCATGATTAAGACTAATTTCACGATAAA |
| 146 | 40 | CCTCGCCGTTGTATCACCGTCACCGAAACGAGGATTACTA |
| 147 | 40 | TCTTTGACAGTAGCAATACCAAGTTACAAATATTACGCCA |
| 148 | 40 | GCCAAAGAAATTATCACCGGAACCAGAGGCTTTAGAACGC |

| | | |
|---|---|---|
| 149 | 40 | CCAGTAATCCTTCCAGTAAGCGTCATAAAAAAGCTTATCC |
| 150 | 39 | CCGCCACCCAAACCAGCAGAAGATATTTGGAAACCGGAA |
| 151 | 39 | TCCAACGCATCGCCACAATGAAATAGCAGACTAATATCA |
| 152 | 37 | GAATGGATTGCTTTATTACCGCGCCCAATATCAAATC |
| 153 | 37 | TGCTCAGAGCATTCTATCCCAATCCAAATATCGATTT |
| 154 | 25 | ACTCCAGCTAATGCGAGGACTAAAG |
| 155 | 25 | GCATCAATAGAAGGGCTCCAAAAGG |
| 156 | 21 | TTTACACGAGATTATAAATCA |
| 157 | 21 | TCGCCATCATTTAAAACATCA |
